# Adaptive motility enables neutrophils to rapidly navigate confined capillaries

**DOI:** 10.1101/2025.01.08.631737

**Authors:** Mathieu Deygas, Mathilde Bernard, Pierre Nivoit, Lucie Barbier, Li Wang, Mathieu Maurin, Emmanuel Tejerina, Theresa Jakuszeit, Alexandre Deslys, Moira Garcia-Gomez, Oumaima Baaziz, Serge Garbay, Emmanuel Terriac, Rafaele Attia, Guillaume Duménil, Matthieu Piel, Pablo Vargas

## Abstract

As the first responders of the immune system, neutrophils rapidly and abundantly reach inflamed tissues through blood capillaries. The diameter of capillaries can be as narrow as two microns, imposing considerable deformations on neutrophils. Notably, capillary obstruction due to neutrophil retention causes vascular dysfunction and contributes to the pathogenesis of several diseases. However, the cellular mechanisms that allow neutrophils to migrate into small capillaries and to avoid retention remain unknown. In this study, we demonstrate, both *in vivo* and *in vitro*, that capillary size does not influence neutrophil migration velocity. During migration into capillaries of different sizes, neutrophils maintain high speed, a phenomenon associated with a global actomyosin cytoskeleton rearrangement in response to confinement strength. In irregular capillaries, neutrophils rapidly adapt their cell contractility via the ROCK-MyoII pathway, which allows them to sustain their migration speed along the vessels despite changes in confinement. At the single cell level, inhibition of ROCK impairs actomyosin cytoskeleton rearrangement and reduces neutrophil migration speed within confined capillaries. At the collective level, ROCK inhibition hampers efficient neutrophil trafficking in a network of small capillaries, resulting in vessel obstruction. These findings reveal a unique capacity of neutrophils to rapidly and dynamically adapt their migration to the confinement strength of capillaries, an ability that might limit vascular dysfunction during inflammation.

## INTRODUCTION

Neutrophils are the most abundant immune cells in blood circulation, serving as the first line of defense against infections. They act as rapid responders to threats, capable of reaching speeds of up to 30 μm/min during inflammation, significantly faster than other cells (*1*). This remarkable speed is partly due to their plasticity, crucial for their fast displacement in various tissue microenvironments (*2*). However, excessive recruitment of neutrophils can lead to organ dysfunction (*3,4*), highlighting the need for tight control of their recruitment, retention and clearance.

Upon tissue injury and infection, circulating neutrophils slow down and crawl on the activated endothelium at the level of post-capillary venules, where they eventually stop and extravasate (*5*). Neutrophils also navigate within networks of blood capillaries whose organization and diameters, ranging from 2 to 15 µm, vary between organs (*6*). Consequently, neutrophils experience different levels of confinement and can undergo substantial deformation within narrow capillaries. In this context, neutrophils migrate, independently of the blood flow, in 3 µm-diameter capillaries in tissues such as the skin and the lymph nodes (*7–9*) (Figure 1a). In such small capillaries, neutrophils become elongated and must counteract the resistance imposed by the confinement strength (*10–13*). However, little is known about the ability of neutrophils to ensure their migration in such microenvironments.

**Figure 1:**
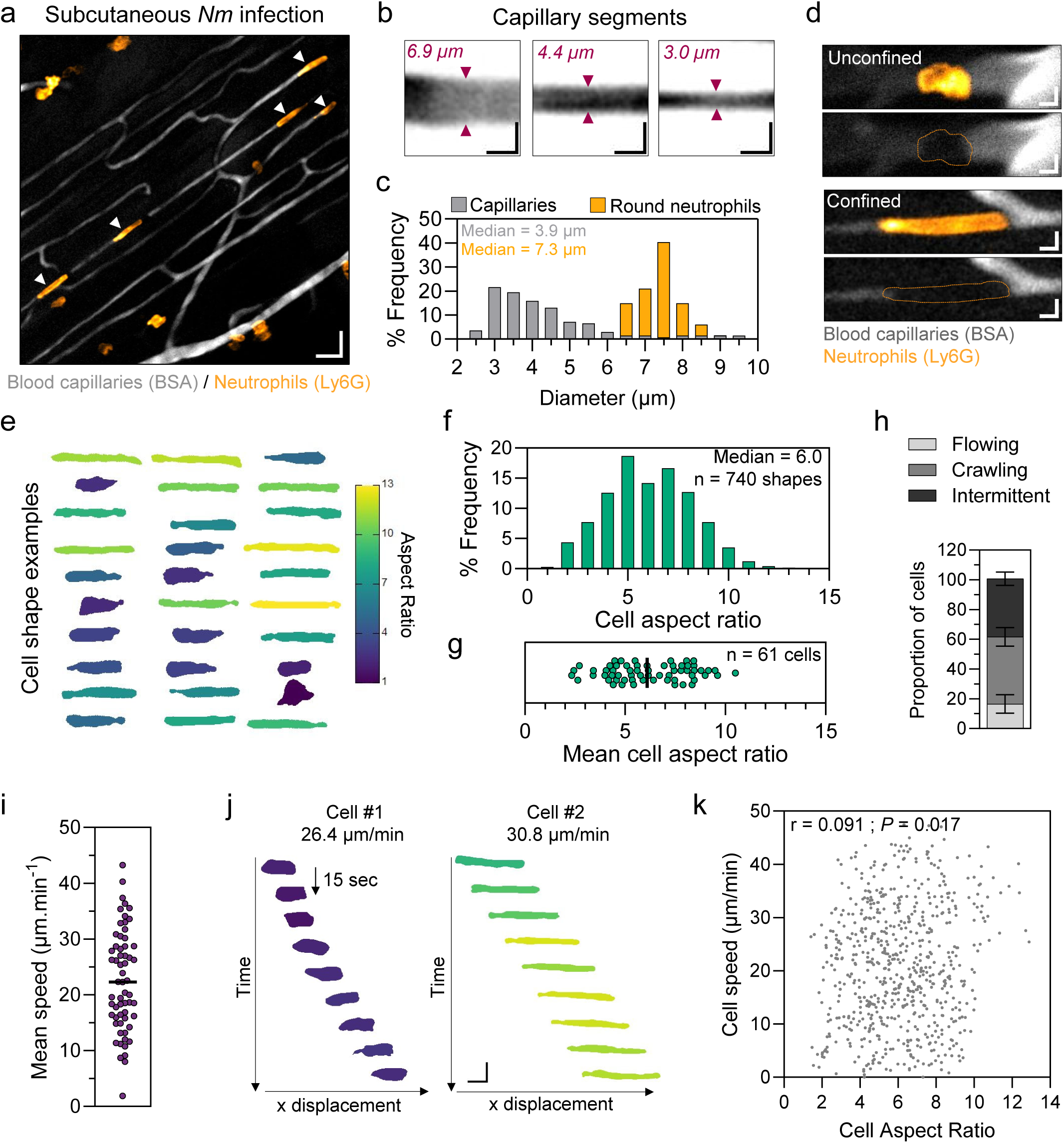
Neutrophil migration in skin capillaries is independent of confinement strength. **a**. Spinning disk-confocal images of subcutaneous blood capillaries (gray, fluorescent BSA) and endogenous neutrophils (orange, Ly6G) two hours after *N.meningitidis* local injection. Neutrophils located inside capillaries are indicated with white arrow. Scale bar: 20 µm. **b**. Magnifications of three capillary segments revealed by fluorescent BSA. Scale bar: 5 µm. **c**. Frequency distribution (percentage) of capillaries and rounded neutrophils (in large vessels) diameters. Data are pooled from three independent experiments, for a total of 139 capillary segments and 67 rounded neutrophils. **d**. Magnification of two cells trafficking inside the microvasculature. Scale bar: 5 µm. e. Montage of 30 cells trafficking in subcutaneous capillaries, pseudo-colored by the value of their AR. **f**. Frequency distribution of cell AR of neutrophils trafficking inside capillaries. n=740 shapes from 3 independent experiments. **g**. Mean cell AR of 61 cells from 3 independent experiments. **h**. Percentage of flowing, crawling and intermittently flowing and crawling cells inside subcutaneous capillaries from 3 independent experiments (total of 229 cells). **i**. Migration speed of individual cells in subcutaneous capillaries, n = 61 cells from 3 independent experiments. **j**. Kymographs (in the laboratory frame of reference) of two neutrophils migrating in subcutaneous capillaries of different diameters. Cell AR is represented in pseudo-color corresponding to the legend in fig 1e. **k**. Scatter plots comparing instantaneous speed and instantaneous cell AR of cells in subcutaneous capillaries. n = 679 timepoints from 61 cells from 3 independent experiments. Spearman correlation coefficient = 0.09, P = 0.017 (two tailed).

Notably, impaired neutrophil migration in confined capillaries can cause vascular obstruction and lead to tissue damage. For instance, in Alzheimer’s disease, ischemic stroke, and epilepsy, the arrest of individual neutrophils in brain capillaries contributes to vessel stalling and promotes vascular dysfunction (*14–17*). Moreover, the intravascular retention of neutrophil in lung capillaries can lead to formation of “trains” of cells, reducing blood flow and impairing oxygenation (*18*).

Microfabricated tools have enabled the identification of mechanisms that neutrophils use to efficiently navigate capillary networks. For example, by sensing hydraulic resistance, neutrophils can avoid accumulation in dead-ends and help their trafficking through complex vessel-like networks (*8,19,20*). Notably, neutrophils and other leukocytes may use their nucleus as a sensor to avoid confined areas with high resistance (*21*). Nonetheless, the cellular mechanisms that allow neutrophils to navigate through capillaries without blocking small vessels remain mostly unexplored.

Here, we investigated neutrophil migration in capillaries both *in vivo* and *in vitro*. We show that neutrophil migration speed is independent of the cell elongation they reach in artificial vessels and *in vivo* capillaries. Neutrophils can also sustain their fast speed in capillaries that impose confinement changes. This capacity results from an active process during which neutrophils dynamically adapt their actomyosin cytoskeleton in response to the confinement strength. Notably, inhibition of actomyosin contractility prevents speed adaptation to confinement and triggers neutrophil jamming in networks of small capillaries. This work highlights a cell intrinsic mechanism setting neutrophil speed in capillary vessels, required for their migration under strong confinement. Failure of this machinery causes neutrophil retention within narrow capillaries, potentially compromising tissue integrity during innate immune responses.

## RESULTS

### Neutrophil migration in skin capillaries is independent of confinement strength

Neutrophils are prone to undergoing significant deformations when they migrate into small blood capillaries. However, the impact of these deformations on their motility remains unknown. To investigate neutrophil migration in confined blood capillaries *in vivo*, we conducted subcutaneous live cell imaging of endogenous neutrophils in mouse flank skin. In this context, neutrophil migration in capillaries can occur independently of blood flow during inflammation, as previously reported (*7*). Imaging of the flank skin capillary network revealed a parallel array of tubes with diameters ranging from 2.5 to 11.2 µm (median = 3.96 µm) (Figure 1a-1c). In comparison, the median diameter of unconfined neutrophils was 7.3 µm (Figure 1c). In the absence of tissue inflammation, neutrophils were barely detected (Supplementary figure 1a, supplementary movie S1). Local injection with *Neisseria meningitidis* (*Nm*) triggered neutrophil accumulation in capillaries, which we observed 1-hour post-treatment (Figure 1a, supplementary figure 1b, 1c). In the next hours, neutrophils accumulated around *Nm* in the tissue (Supplementary figure 1c-e, supplementary movie S2). Notably, no extravasation was observed at the level of the capillaries (Supplementary movie S3). These results show neutrophil migration through micrometric capillaries at early stages of inflammation in the skin in mice flanks.

Morphometric analysis of neutrophils within capillaries 1-hour post-infection revealed a wide range of confinement-induced cell deformations, as measured by the cell aspect ratio (AR, cell length divided by cell width) (Figure 1d-1g). While some neutrophils were almost round, others were ten times longer than wide, with an average AR of 6 (Figure 1e-1g, Supplementary figure 1f). Analysis of neutrophil migration in capillaries using live-cell imaging showed that 40% of cells were crawling (speed < 50 µm/min, median speed 22 µm/min), 20% were flowing (speed > 50 µm/min), and 40% alternated between flowing and crawling phases (Figure 1h, i, Supplementary figure 1g-i). We observed some neutrophils changing direction within the tubes or migrating against blood flow, which reflected their active migration (Supplementary figure 1j, supplementary movie S4). Strikingly, the instantaneous speed of neutrophils did not depend on the extent of cell elongation (Figure 1j, k, supplementary movie S3). This finding shows that the migration of endogenous neutrophils in blood vessels inducing high cell elongations is not hampered by the confinement imposed by the capillary tube in this *in vivo* system.

### Neutrophil migration is independent of confinement strength into *in vitro* capillaries

While *in vivo* results provide valuable insights on neutrophil dynamics in capillaries, they cannot fully exclude the potential influence of microenvironmental factors at the site of inflammation. Adhesion molecules are modulated by the inflammatory state of the tissue and might contribute to regulating neutrophil migration *in vivo*. In addition, blood flow can modulate neutrophil migration in capillaries. To isolate the effect of physical constraints on neutrophil migration, we employed microfabrication to create *in vitro* capillaries with varying cross-section areas (ranging from 3 to 8 µm in width and 4 µm in height), mimicking the diameter range of the flank skin vessels observed *in vivo* (Figure 2a). As a cellular model, we used primary bone-marrow neutrophils matured by treatment with Granulocyte-Macrophage Colony Stimulating Factor (GM-CSF) (Supplementary figure 8) (*22–24*). Mature mouse neutrophils (mNeu) entered spontaneously into fibronectin-coated microchannels of all dimensions tested, in the absence of any chemoattractant (Figure 2a-b, Supplementary figure 2a). Morphometric analysis of mNeu migrating in the *in vitro* capillaries showed a correlation between cell elongation and confinement strength. The average cell AR ranged from 2.3 in 8 µm-width microchannels to 9.3 in 3 µm-width microchannels (Figure 2c, supplementary figure 2c). These values fall within the range of *in vivo* measurements (Figure 1f). The nucleus AR also increased as a function of the confinement strength, indicating complete cell deformation (Supplementary figure 2b). Single-cell migration analysis of mNeu revealed an equivalent average speed across all tested dimensions (around 17 µm/min) (Figure 2d, supplementary figure 2d). Similarly to *in vivo* data, cell migration speed in microchannels of different sizes remained independent of the cell AR (Figure 2e, supplementary movie S5). These findings demonstrate that spontaneous mNeu migration is not significantly affected by the confinement strength imposed by the *in vitro* capillaries.

**Figure 2:**
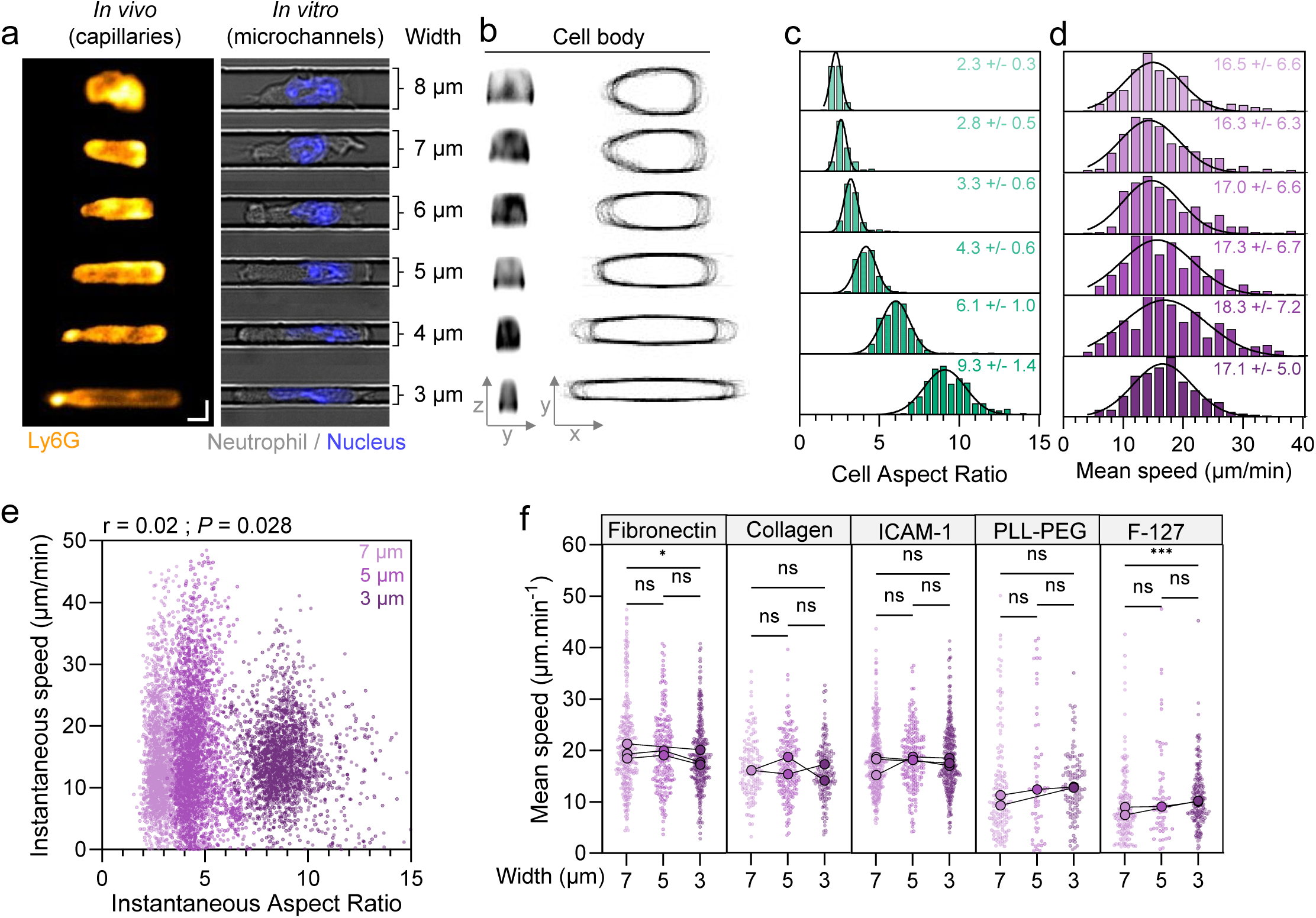
Neutrophil migration is independent of confinement strength into *in vitro* capillaries. **a**: Examples of in vivo neutrophils in subcutaneous capillaries (left) and in vitro bone-marrow neutrophils in microchannels of different widths (right). Scale bar 5 µm. **b**: Cross sections of LifeAct-EGFP expressing neutrophils (left) and representation of 30 cell contours (sµm) per channel width (top view, right). **c**: Distribution of mean cell AR, data pooled from 3 independent experiments. Median +/- SD is indicated for each channel width. 8 µm, n= 183; 7 µm, n = 150; 6 µm, n = 205; 5 µm, n = 218; 4 µm, n = 231; 3 µm, n = 214. **d**: Distribution of mean cell speed. Data are pooled from 5 independent experiments. Median +/- SD is indicated for each channel width. 8µm, n = 289; 7µm, n = 263; 6 µm, n = 333; 5 µm, n = 314; 4 µm, n = 345, 3 µm, n = 350. **e**: Cell AR correlation to cell speed of neutrophils. Dots, individual cell shape/speed value of neutrophils in 7, 5 and 3 µm channel width. Data are representative of 3 independent experiments. 7 µm, n = 2189; 5 µm, n = 2876; 3 µm, n = 2137. Spearman correlation coefficient = 0.02, P = 0.056 (two-tailed). **f**: Mean cell speeds of neutrophils migrating in microchannels of 7, 5, 3 µm widths coated with different surface molecules, from 2 or 3 independent experiments. Median of each experiment are represented by paired bigger dots. Fibronectin 10 µg/ml, 7 µm, n = 277; 5 µm, n = 209; 3 µm, n = 289. Collagen 10 µg/ml, 7 µm, n = 134; 5 µm, n = 180; 3 µm, n = 163. ICAM-1 10 µg/ml, 7 µm, n = 303; 5 µm, n = 187; 3 µm, n = 349. PLL-g-PEG 1 mg/ml, 7 µm, n = 145; 5 µm, n = 55; 3 µm, n = 117. Pluronic F-127 5% (w/v), 7 µm, n = 167; 5 µm, n = 62; 3 µm, n = 181. Kruskal-Wallis test, ns not significant, * P < 0.05, *** P < 0.001.

Next, we investigated the potential role of specific cell adhesion molecules in influencing the independence of mNeu migration speed from confinement strength. To this end, we repeated the experiment using *in vitro* capillaries coated with either collagen or Intercellular Adhesion Molecule-1 (ICAM-1). Consistent with our previous observations with fibronectin coating, mNeu exhibited similar average speeds at different confinement strengths in both conditions (Figure 2f). These results showed that neutrophil migration speed in microfabricated vessels of varying sizes was independent of the adhesive substrates tested.

To reduce the contribution of specific cell adhesion molecules, we passivated the microchannel surfaces with PLL-PEG or Pluronic F-127. In these conditions, average speed was nearly 50 % slower compared to conditions with adhesive molecules present (Figure 2f) but remained unaffected by the confinement strength (Figure 2f). These findings show mNeu can maintain their migration speed at different confinements strengths within *in vitro* capillaries in a cell adhesion-independent manner.

### Neutrophils adapt their migration to changes in confinement strength

We next explored whether neutrophil migration could be affected by changes in confinement strength, a scenario encountered *in vivo* (Supplementary figure 3a, supplementary movie S6). To investigate this, we first assessed mNeu migration in artificial capillaries featuring a sharp decrease in their width from 6 µm to 3 µm (maintaining a constant height of 4 µm) (Figure 3a). As expected, mNeu entry into the smaller vessel section led to an increase in cell AR (Figure 3a, b and Supplementary figure 3b). Notably, despite significant cell elongation, mNeu showed equivalent average speeds in both the wide and narrow sections of the capillary (Figure 3b-3c, supplementary movie S7). A transient decrease in cell speed was observed during the cell transit from the larger to the smaller section (Figure 3b, supplementary figure 3b). This indicated that these cells are sensitive to changes in confinement strength but can rapidly recover their migration speed. Single-cell analysis revealed that the speed ratio between the two sections was close to one, demonstrating that individual mNeu maintained their average speed across different confinement strengths (Figure 3d). Approximately 50% of the cells navigated the transition in less than 2 min (Figure 3e and Supplementary figure 3c) and 95% passed in less than 6 min (Supplementary figure 3c).

**Figure 3:**
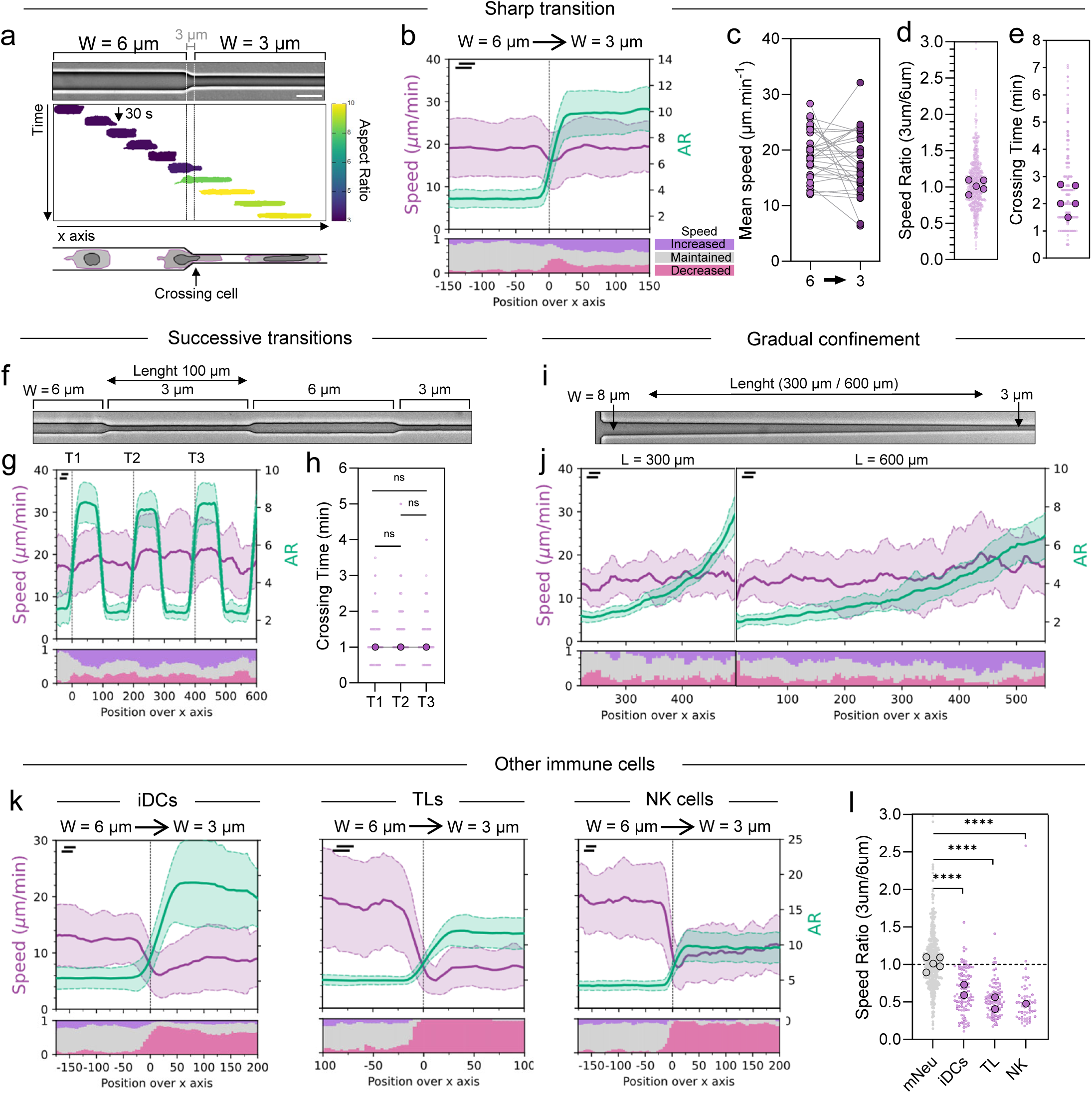
Neutrophils adapt their migration to changes in confinement strength. **a**: Migration of a neutrophil, pseudo-colored by its aspect ratio, in a size-transition microchannel harboring a sharp narrowing from 6 µm to 3 µm width (4 µm height). **b**: Top, evolution of cell speed and cell aspect ratio over the x position in a size-transition microchannel. Position 0 corresponds to the necking. Bottom, evolution of the fraction of cells displaying a speed increase, maintenance or decrease over the x axis, compared with the respective mean speed in the 6 µm wide section. Data are one representative of 5 experiments, N = 122 cells. **c**: Average speeds of 30 cells migrating from 6 µm to 3 µm wide section. **d**: Speed ratio: average speed in the 3 µm wide section normalized by the average speed in the 6 µm wide section. Each small dot represents a cell, and bigger dots represent a median of one experiment. From 5 independent experiments, N = 544 cells. **e**: Crossing time or time spent in the necking (see bottom drawing figure 3a). Each small dot represents a cell, and bigger dots represent a median of one experiment. From five independent experiment, n = 544 cells. **f**: Image of a microchannel with successive transition between 6 µm and 3 µm widths. **g**: Migratory behavior of neutrophils in a successive size-transition microchannel. Top, evolution of cell speed and cell aspect ratio over the x position in a successive size-transition microchannel. Transitions (T1, T2, T3) from 6 µm to 3 µm are indicated with dotted vertical lines. Bottom, evolution of the fraction of cells displaying a speed increase, maintenance or decrease over the x axis, compared with the respective mean speed in the first 6 µm wide section. Data are one representative of 2 experiments, n = 90 cells. **h**: Crossing times for three consecutive transitions (6 µm to 3 µm wide section). Dots, individual cells. Data are one representative of 2 experiments, from 2 technical replicates, T1, N = 128 cells, T2, N = 139 cells, T3, N = 109 cells. **i**: Image of a funnel-shaped microchannel with gradual confinement from 8 µm to 3 µm width. **j**: Migratory behavior of neutrophils in a funnel-shaped microchannel, transitioning gradually from 8 to 3 µm width. Top, evolution of cell speed and cell aspect ratio over the x position in a funnel-shaped microchannel of 300 µm length (left) or 600 µm length (right). Bottom, evolution of the fraction of cells displaying a speed increase, maintenance or decrease over the x axis. Data are one representative of 2 and 3 experiments (300 and 600 µm length respectively). L = 300 µm, n = 21 cells. L = 600 µm, n = 23 cells. **k**: Migratory behaviors of immature dendritic cells (iDCs), T-Lymphocytes (TL) and natural killer cells (NK) in a size-transition microchannel. Top, evolution of cell speed and cell aspect ratio over the x position in a size-transition microchannel. Position 0 corresponds to the necking. Bottom, evolution of the fraction of cells displaying a speed increase, maintenance or decrease over the x axis, compared with the respective mean speed in the 6 µm wide section. Data are one representative of 2 (iDC), 2 (TL) and 1 (NK) experiments, iDCs, n = 79 cells; TL, n = 38 cells; NK, n = 56 cells. **l**: Speed ratio of iDC, TL and NK cells in size-transition microchannels. Dots, individual cells. iDCs, n = 103 cells; TL, n = 96 cells; NK, n = 58 cells. In **b**, **g**, **j**, **k** bins size is indicated by the bar length and the sliding window is indicated by the shift between the two bars (upper left).

Similar experiments where mNeu migrated from 3 µm to 6 µm width vessels also showed speed maintenance (Supplementary figure 3d-h), with all cells completing the transition in less than 5 min (Supplementary figure 3h). We then challenged neutrophil migration in capillaries alternating successive 6 µm and 3 µm wide sections (Figure 3f, supplementary movie S8). Overall, average mNeu speed was maintained among the different sections of this setup (Figure 3g, supplementary figure 3i). Notably, single-cell analysis showed that the crossing time for passing between adjacent sections was not altered (Figure 3h). These results demonstrate that neutrophils can maintain their average migration speed in capillaries with alternating confinement changes *in vitro*, highlighting their adaptive motility in response to varying confinement strengths.

To further investigate neutrophil migration in irregular microenvironments, we assessed mNeu migration in funnel-shaped capillaries decreasing their width from 8 µm to 2 µm over a distance of either 300 µm or 600 µm (Figure 3i, supplementary movie S9). Morphometric analysis showed gradual increase in cell AR as mNeu migrated from the wide to the narrow side of the funnel (Figure 3j and supplementary Figure 3j). Importantly, in both cases, mNeu speed remained constant despite increased confinement strength (Figure 3j and supplementary Figure 3j). This data demonstrates that neutrophils can maintain their migration speed in funnel-shaped capillaries *in vitro*, highlighting their ability to navigate and adapt to varying degrees of confinement.

To determine if the ability to maintain migration speed in confining capillaries was specific to neutrophils, we examined the motility of other primary leukocytes in capillaries with a sharp narrowing from 6 µm to 3 µm in width. Analysis of immature dendritic cells (iDCs), T lymphocytes (TLs) and Natural killer (NK) cells revealed a decrease in migration speed upon passage into the confined vessel section (Figure 3k). Single-cell speed ratio across both capillary sections showed a reduction of 65%, 50% and 50% for iDCs, TLs and NKs, respectively (Figure 3l). These findings demonstrate that primary neutrophils possess a cell-intrinsic capacity to maintain their migration speed upon abrupt or gradual changes in the confinement strength imposed by a capillary tube *in vitro*. This property appears to be specific to neutrophils, as other leukocytes failed to maintain their migration speed in capillaries imposing increased confinement strength.

### Neutrophil F-actin polarity is regulated by confinement strength

Previous reports have shown that cells might rapidly adapt their cytoskeleton in response to confinement (*25–29*). Therefore, we hypothesized fast F-actin remodeling in mNeu could be associated with speed maintenance in response to acute changes in confinement strength.

To address F-actin distribution under confinement, we performed spinning-disk live-cell imaging of Lifeact-EGFP expressing mNeu migrating in microfabricated capillaries imposing a sharp width reduction. In the wider section of the vessels, F-actin was preferentially located at the cell front, as previously described for neutrophil-like cells (*30*). Notably, passage to the narrower section of the capillary was accompanied by a fast F-actin shift to the cell rear (Figure 4a-c and Supplementary figure 4a-c, supplementary movie S10). In both sections, F-actin was found to be mostly cortical, despite its differential localization in cells (Figure 4a-b, supplementary figure 4b).

**Figure 4:**
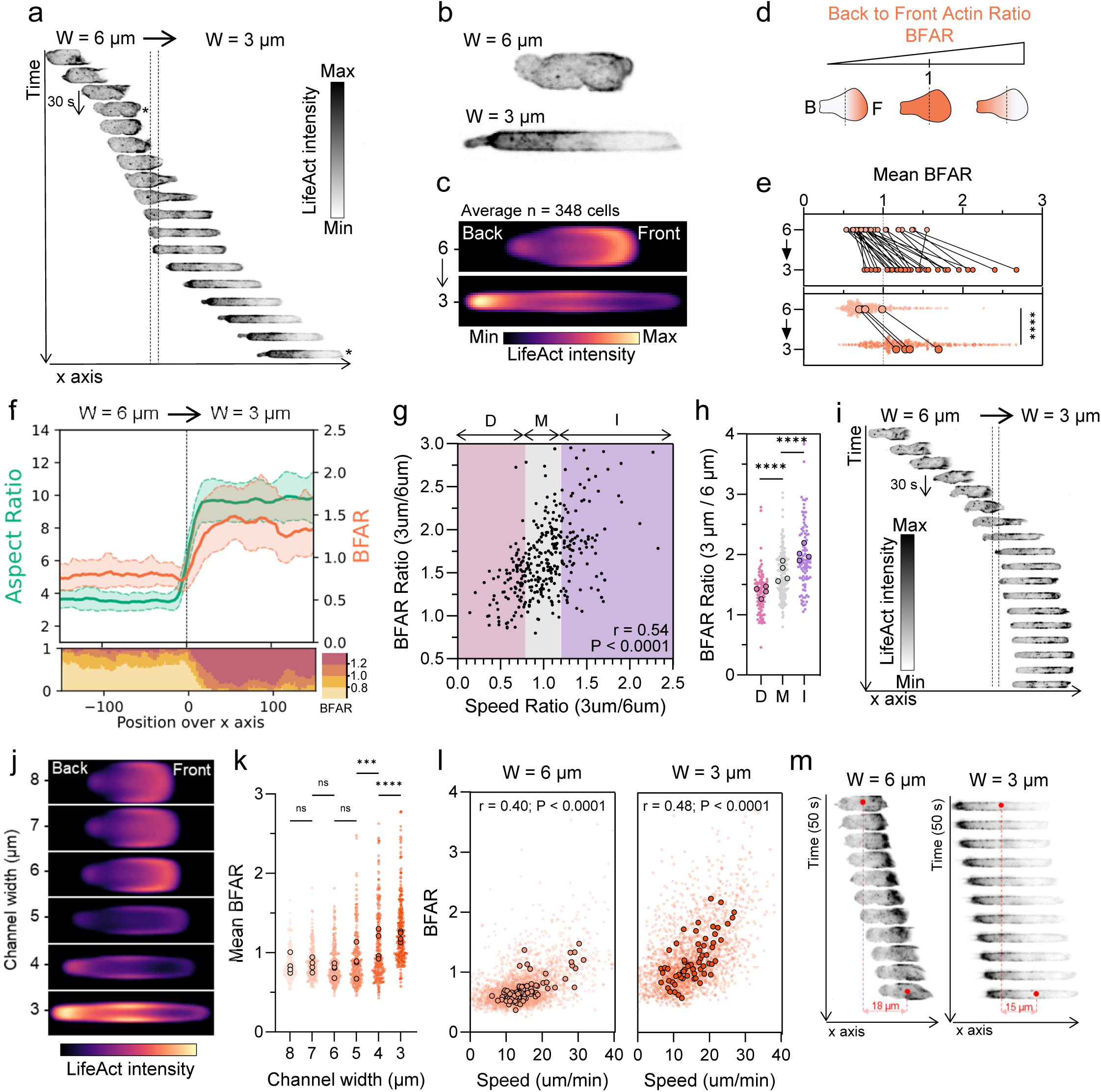
Neutrophil F-actin polarity is regulated by confinement strength. **a**: Kimograph of a Lifeact-EGFP expressing neutrophil migrating in size-transition microchannel. Each row represents a timepoint (30 seconds). Lifeact-EGFP intensity is represented in inverted gray color. **b**: Magnification of the cell at timepoints indicated by * in a. The cell is migrating toward the right in 6 µm wide (top) and 3 µm wide section (bottom). **c**: Average Lifeact-EGFP intensity of n = 348 cells migrating from 6 µm to 3 µm wide section. Data pooled from 4 independent experiments. **d**: Schematic representation of Back to Front Actin Ratio (BFAR). **e**: Top, mean BFAR of n = 30 individual neutrophils migrating from the 6 µm to the 3 µm wide section. Bottom, mean BFAR of neutrophils migrating from the 6 µm to the 3 µm wide section. Light dots, individual cells, n = 348 cells. Dark dots, BFAR medians of 4 independent experiments. Wilcoxon test (paired), **** P < 0.0001. **f**: Top, BFAR and cell AR over the x position in a size-transition microchannel. Position 0 corresponds to the necking. Bottom, fraction of cells over the x axis, with BFAR values between indicated BFAR ranges. Data are one representative of 4 experiments, n = 50 cells. **g**: Scatter plot showing the correlation between BFAR ratio and speed ratio. Ratios are defined by the mean values in 3 µm section divided by mean values of 6 µm section for individual cells. Speed decrease (D), maintenance (M) and increase (I) are indicated with colored rectangles. Data pooled from 4 independent experiments, n = 346 cells. Spearman correlation coefficient = 0.54, P < 0.0001. **h**: BFAR ratios of cells depending on their migratory response to sharp change in confinement strength. Light dots, individual cells; dark dots, mean of experiment. Speed decrease (D), n = 90; Speed maintenance (M), n = 157; Speed increase (I), n = 99. Unpaired Mann Whitney test, **** P < 0.0001. **i**: Kimograph of a Lifeact-EGFP expressing neutrophil slowing down in size-transition microchannel. Each row represents a timepoint (30 seconds). Lifeact-EGFP intensity is represented in inverted gray color. **j**: Average Lifeact-EGFP intensity mapping of migrating neutrophils in straight microchannels of different widths. Data are 60 cells per channel width, from 3 independent experiments (20 cells per experiment). **k**: Kimographs of Lifeact-EGFP expressing neutrophils migrating in 6 µm wide (left) and 3 µm wide straight microchannels for 50 seconds. Lifeact-EGFP intensity is represented in inverted gray color. **l**: Mean BFAR of migrating neutrophils in straight microchannels of different widths. Data are pooled from 4 independent experiments. Light dots, individual cells; dark dots, median of experiment. 8 µm, n = 216; 7 µm, n = 210; 6 µm, n = 256; 5 µm, n = 289; 4 µm, n = 271, 3 µm, n = 253. Kruskall-Wallis test. ns = not significant; *** P < 0.001; **** P < 0.0001. **m**: Relation between BFAR and cell speed of neutrophils migrating in 6 µm and 3 µm wide straight microchannels. Light dots, instantaneous timepoint; dark dots, mean of individual cells. Data are one representative of 4 independent experiments. Spearman correlation coefficient, 6 µm, r = 0.40; 3 µm, r = 0.48, P < 0.0001.

The front-to-back F-actin shift was confirmed at the population level by computing the average F-actin signal in both sections of the capillaries (Figure 4c, supplementary figure 4c). These results highlight a differential F-actin organization in mNeu migrating at diverse confinement strengths. To get insights into F-actin dynamics at the single-cell level, we used the back-to-front actin ratio (BFAR) as a metric (Figure 4d). On average, this indicator was lower than 1 in the 6 µm-width vessel section, indicating preferential F-actin localization at the cell front (Figure 4e). Upon cell passage into the narrower section, the average value increased above 1, indicating a front-to-back switch (Figure 4e). Spatial analysis showed that the BFAR increased concomitantly with mNeu elongation (Figure 4f and Supplementary figure 4d). Of note, single-cell analysis showed a positive correlation between the BFAR and the speed ratio at each time step (Figure 4g, h). This result indicates that the migratory behavior of neutrophils in the size-transition channel is directly associated with the redistribution of F-actin at the cell rear. This observation is exemplified by cells incapable of concentrating F-actin at their back, which failed to adapt their speed in response to an increase in confinement strength (Figure 4i, supplementary movie S10). Of note, when mNeu migrated from the narrow to the wider section of the capillaries, F-actin quickly re-localized to the cell front (Supplementary figure 4e-g, supplementary movie S10). These results show rapid and reversible F-actin reorganization in response to confinement strength during neutrophil migration in capillaries.

To evaluate the sensitivity of the F-actin rearrangements in response to the confinement strength, we calculated the average F-actin signal in mNeu migrating into capillaries of different widths (Figure 2a). In the range of 8 µm to 6 µm width, average F-actin signal was mostly found at the cell front (Figure 4j, k). In the range from 5 µm to 3 µm width capillaries, F-actin gradually accumulated at the cell rear as cells became more elongated (Figure 4j, k, supplementary movie S11). The fraction of neutrophils with an average F-actin signal at the rear also increased with confinement (Supplementary figure 4h, i). These results indicate that F-actin enrichment at the cell rear in mNeu is dependent on the confinement strength.

Notably, speed and BFAR showed a positive correlation at different confinement strengths, both at the instantaneous and cell-average level (Figure 4l and Supplementary figure 4j). This result indicates that faster cells display more rear F-actin than slower cells, for any confinement strength. More generally, when considering cells migrating at a given speed at different confinements, rear F-actin was stronger in narrower vessels (Figure 4m, supplementary figure 4j, supplementary movie S11). Altogether, these data demonstrates that mNeu can rearrange their actin cytoskeleton in the scale of seconds in response to changes in the confinement strength during their migration in capillaries *in vitro*. It further suggests that maintaining a given speed in narrower channels requires increased rear F-actin in these cells.

### Myosin II is required for neutrophil migration adaptation to confinement strength

The main driver for F-actin contraction and cell migration in leukocytes is the molecular motor non-muscle Myosin IIA (MyoII) (*31*). Therefore, we investigated whether MyoII activity could be regulated by confinement strength in capillaries *in vitro*. For that, we performed immunofluorescence of the phosphorylated Myosin Light Chain 2 (pMLC2), an indicator of MyoII activation. In neutrophils migrating from 6 µm to 3 µm width capillaries, pMLC2 signal was higher in the narrower vessel section, indicating increased MyoII activation under confinement (Figure 5a, Supplementary figure 5a-c).

**Figure 5:**
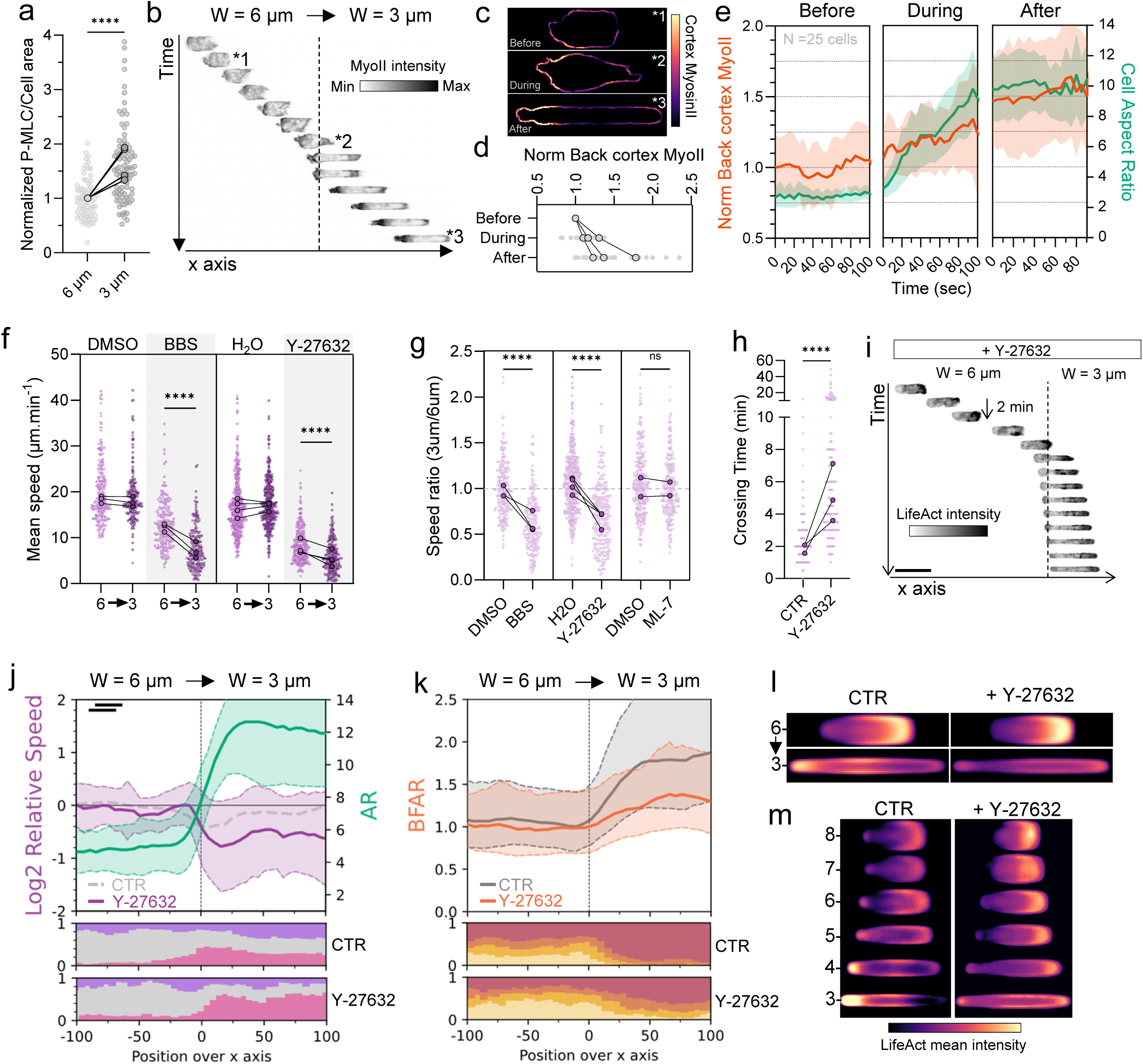
Myosin II is required for neutrophil migration adaptation to confinement strength. **a**: P-MLC2 levels per cell area. Data are normalized by the average P-MLC2 level in the 6 µm-wide microchannel of each experiment. Data are pooled from 4 independent experiments; 6 µm, n = 73 cells; 3 µm, n = 91 cells. Mann Whitney test, **** P < 0.0001. **b**: Kimograph of a MyoII-GFP expressing neutrophil migrating in size-transition microchannel. Data represent the signal in the middle cross-section of the cell obtained with spinning disk confocal microscope. **c**: Cortical MyoII-GFP intensity of 3 timepoints (before, during and after the narrowing) indicated in **b**. **d**: Normalized cortical MyoII intensity (of the 25 % cell back) before, during and after the passage of mNeu through the narrowing. Data are pooled from 3 independent experiments, n = 25 cells. Median of each experiment is represented with connected big dots. **e**: Evolution of cortical MyosinIIA-GFP intensity (25 % back of the cell) and cell aspect ratio over time of mNeu migrating in a size-transition microchannel. Data are pooled from 3 independent experiments, n = 25 cells. Time windows are separated according to the position of cells: before, during and after the passage through the narrowing. **f**: Average speeds of cells migrating from 6 µm to 3 µm wide section, treated or not with Blebbistatin (25 µM) or Y-27632 (25 µM). Data are pooled from 3 (BBS) and 4 (Y-27632) independent experiments. DMSO, n = 196 cells; BBS, n =188 cells; H20, n = 376 cells, Y-27632, n = 231 cells. Mann Whitney test, P < 0.0001. **g**: Speed ratio of neutrophils treated or not with BBS (25 µM), Y-27632 (25 µM) or ML-7 (20 µM). Each light dot represents a cell, and dark dot represents a median of one experiment. Data are pooled from 3 (BBS), 4 (Y-27632) and 2 (ML-7) independent experiments. DMSO, n = 196 cells; BBS, n =188 cells; H20, n = 376 cells, Y-27632, n = 231 cells. DMSO (ML-7), n = 257; ML-7, n = 217. Mann Whitney test; ns, not significant; **** P < 0.0001. **h**: Crossing time of neutrophils treated or not with Y-27632 (25 µM). Data pooled from 3 independent experiments, CTR, n = 363; Y-27632, n = 146 cells. **i**: Kimograph of a Lifeact-EGFP expressing neutrophil treated with Y-27632 (25 µM) migrating in size-transition microchannel. Each row represents a timepoint (2 minutes). Lifeact-EGFP intensity is represented in inverted gray color. **j**: Top, speed and aspect ratio of Y-27632 treated neutrophils over the x position in a size-transition microchannel. Speed of the control condition is indicated in gray dotted line. Position 0 corresponds to the necking. Bottom, fraction of control and Y-27632 cells over the x axis, displaying a speed increase, maintenance or decrease over the x axis, compared with the respective mean speed in the 6 µm wide section. Data are one representative of 3 experiments, n = 95 cells. **k**: Top, BFAR of Y-27632 treated neutrophils (orange) and control cells (gray) over the x position in a size-transition microchannel. Bottom, fraction of control and Y-27632 cells over the x axis, with BFAR values between indicated BFAR ranges Data are one representative of 3 experiments, n = 95 cells. **l**: Average Lifeact-EGFP intensity control and Y-27632 treated cells migrating from 6 µm to 3 µm-wide section. Data pooled from 3 independent experiments. Control, n = 350 cells; Y-27632, n = 93 cells. **m**: Average Lifeact-EGFP intensity mapping of control and Y-27632 treated cells in straight microchannels of different widths. Data are 40 cells per channel width, from 2 independent experiments.

We then analyzed MyoII dynamics taking advantage of primary neutrophils obtained from MyoII-GFP mice (*32*). Spinning disk live-cell imaging showed MyoII-GFP distribution all over the cell, with cortical puncta and diffuse cytoplasmic sub-localizations (Supplementary figure 5d, e). As previously demonstrated (*33*), MyoII accumulated at the cell rear a few seconds after the initiation of a protrusion during cell reorientation in 2D (Supplementary figure 5d, supplementary movie S12). In addition, cortical MyoII enrichment at the cell rear correlated with mNeu speed in straight microchannels (Supplementary figure 5e-h, supplementary movie S13). These results confirm that cortical MyoII-GFP in mNeu is a proxy for its contractile activity (*34–36*). Notably, cell entry in the 3 µm width vessel section resulted in gradual accumulation of cortical MyoII-GFP at the cell rear (Figure 5b-d, supplementary movie S14). This enrichment reached a 50% increase upon full cell elongation in the narrow microchannel (Figure 5e). These data show that mNeu respond to the confinement imposed by small capillaries through MyoII activation at the cell rear.

To investigate the role of MyoII activity in neutrophil migration adaptation to confinement strength we took advantage of well described pharmacological compounds. Inhibition of the ATPase activity of MyoII using blebbistatin (BBS) decreased average mNeu migration speed in both capillary sections. However, the effect was more prominent on the more confined vessel segment (Figure 5f). In agreement, single-cell analysis of the speed ratio between both sections showed a decrease upon BBS treatment, indicative of defective mNeu migration adaptation to confinement strength (Figure 5g). Similar results were observed upon Rho kinase (ROCK) inhibition (Y-27632), one of the pathways responsible for MyoII activation (Figure 5f-g, supplementary movie S15) (*37*). Consistently, the time taken by mNeu to pass to the confined section of the capillary increased upon ROCK inhibition (Figure 5h-i). However, no effect was detected upon MLC kinase (MLCK) inhibition (ML-7), another pathway which can regulate MyoII activity (Figure 5g and Supplementary figure 5i). This indicates that the ROCK pathway controls mNeu adaptation to confinement strength in capillaries.

Calcium signaling has been shown to be a second messenger key for mechano-responses (*38*). Therefore, we investigated whether calcium dynamics could drive neutrophil migration adaptation to confinement strength. Strikingly, neutrophil passage from the loose to the confined section of capillaries triggered an increase in calcium pulsations (Supplementary figure 5j-l, supplementary movie S16). Surprisingly, extracellular calcium chelation (EGTA) did not affect mNeu ability to adapt their migration to the confinement strength, despite lowering calcium spikes (Supplementary figure 5l-n). In addition, inhibition of mechanosensitive transient receptor potential (TRP) channels (ACA), cationic mechanosensitive channels (GsMTx4), including Piezo, or treatment with a stretch-activated calcium channel blocker (GdCl) and intracellular calcium chelation (BAPTA-AM) did not affect their migration adaptation to confinement (Supplementary figure 5o, p). This data shows that adaptation to changes in confinement strength in neutrophils occurs through a MyoII-dependent cell contractility pathway activated independently of calcium signaling.

To further investigate the role of ROCK in neutrophil migration adaptation to confinement strength, we studied the evolution of mNeu relative speed when facing strong confinement. The results showed that ROCK inhibited cells slowed down as compared to control cells during their passage from the 6 µm to 3µm width capillary sections (Figure 5i-j). ROCK inhibition also affected F-actin dynamics, reducing its accumulation at the cell rear upon passage in the confined vessel section (Figure 5i,k, supplementary movie S16). This result was confirmed by computing the average F-actin signal in both segments of the microchannel, in which ROCK inhibition lowered F-actin signal at the cell rear (Figure 5l). Furthermore, analysis of F-actin distribution in response to confinement using capillaries of different sizes showed that ROCK inhibition prevented F-actin enrichment at the rear as the confinement strength increased (Figure 5m, supplementary figure 5q, r). These results highlight the ROCK signaling pathway as the main driver for neutrophil migration adaptation to confinement.

### Neutrophils adapt cell migration and actin cytoskeleton to confinement *in vivo*

Our *in vitro* results demonstrate that rapid and dynamic actomyosin remodeling allows neutrophils to maintain their fast migration speed in capillaries with varying confinement strengths. To investigate whether similar F-actin rearrangements could also occur during neutrophil migration in capillaries *in vivo*, we transferred LifeAct-EGFP expressing neutrophils into wild type recipient mice and induced local inflammation by injecting *Nm*. Intravital spinning-disk live cell imaging of the subcutaneous capillaries draining the inflamed area showed mostly elongated neutrophils, as expected based on the small size of the capillaries in this tissue. In these cells, the LifeAct-EGFP signal was preferentially located the cell rear (Figure 6a-b, supplementary movie S17). This F-actin distribution was comparable to that observed in 4 and 3 µm width microfabricated capillaries, where cells exhibited a similar AR to those in skin capillaries (Figure 1f-g, and 2c). This data shows that F-actin in neutrophils migrating in small capillaries is similar *in vitro* and *in vivo*.

**Figure 6:**
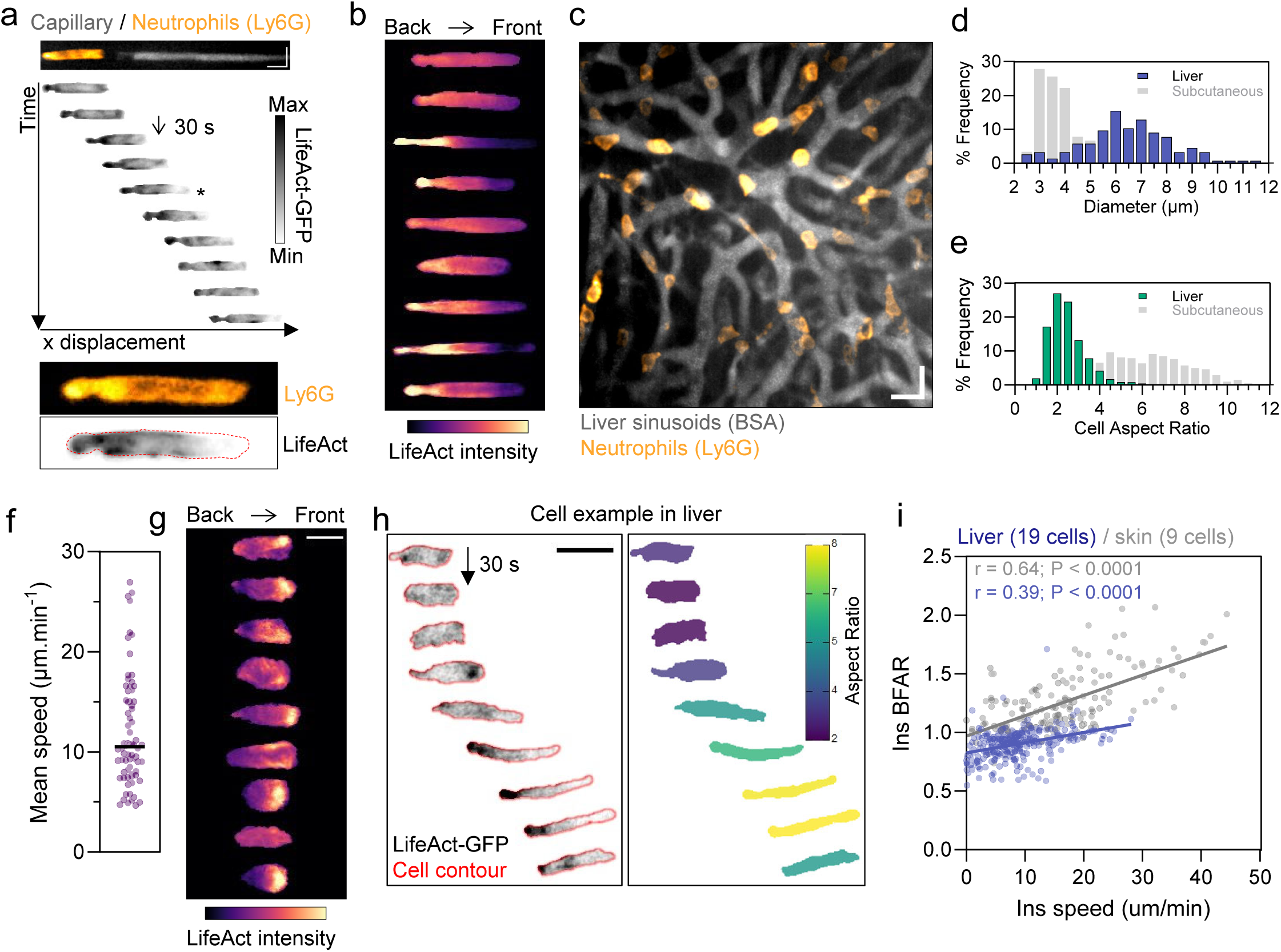
Neutrophils adapt cell migration and actin cytoskeleton to confinement *in vivo*. **a**: Kimograph of a Lifeact-EGFP expressing (stained with anti-Ly6G) neutrophil migrating in subcutaneous capillary. Each row represents a timepoint (30 seconds). Lifeact-EGFP intensity is represented in inverted gray color. Bottom, magnification of the timepoint indicated by *, cell contour in red. **b**: Average Lifeact-EGFP intensity mapping of migrating neutrophils in subcutaneous capillaries. Data are n = 9 cells, from one experiment. **c**: Spinning disk-confocal images of liver sinusoids (gray, fluorescent BSA) and endogenous neutrophils (orange, Ly6G) 2 hours after LPS endotoxemia. Scale bar: 20 µm. **d**: Frequency distribution (percentage) of liver sinusoid diameters. Data are pooled from 4 independent experiments, for a total of 156 segments. **e**: Frequency distribution of cell aspect ratio of neutrophils trafficking inside sinusoids. n=1119 shapes from 3 independent experiments. **f**: Migration mean speed of individual crawling cells in liver sinusoids, n = 72 cells from 3 independent experiments. **g**: Average Lifeact-EGFP intensity mapping of nine migrating neutrophils in liver sinudoids. **h**: Left, kimograph of a Lifeact-EGFP expressing neutrophil migrating in liver sinusoid with a size-transition. Each row represents a timepoint (30 seconds), cell contour in red. Right, kymograph of the same cell color-coded by aspect ratio value. **i**: Scatter plot of the relation between instantaneous BFAR and instantaneous cell speed of n = 174 timepoints from 9 cells (subcutaneous capillaries) and n = 313 timepoints from 19 cells (liver sinusoids). Spearman correlation coefficient (liver) = 0.39, P < 0.0001. Spearman correlation coefficient (skin) = 0.64, P < 0.0001.

We further investigated F-actin dynamics *in vivo* in the liver sinusoids upon LPS endotoxemia (Figure 6c, supplementary figure a, b, supplementary movie S18). In this system, capillaries exhibited higher average diameter and broader distribution compared to skin capillaries (Figure 6d, supplementary figure 6c) (*39*). As a result, neutrophils shown a lower cell AR as compared to skin capillaries (Figure 6e, supplementary Figure 6d, e). The average neutrophil speed in the sinusoids was approximately 10 µm per min, which was lower than in the skin capillaries (Figure 6f, supplementary Figure 6f, g). In these vessels, F-actin was detected primarily at the cell front (Figure 6g, supplementary Figure 6h, supplementary movie S19). This distribution was equivalent to the F-actin pattern observed in 6 µm width microchannels, where neutrophils exhibited a similar cell AR as computed in the liver sinusoids (Figure 2c and 6e). These data show that in mNeu, F-actin organization in capillaries is dictated by their diameter *in vitro* and *in vivo*.

In the liver sinusoids, we detected neutrophils migrating from wider to narrower sections of individual capillaries, although these events were rare. In such situations, similar to our *in vitro* data, we observed a rapid front-to-back F-Actin shift when the mNeu migrated from the loose to the confined areas of the capillary (Figure 6h, supplementary movie S20). This result indicates that neutrophils can rapidly adapt their cytoskeleton organization in response to confinement strength *in vivo*.

We further correlated F-Actin positioning with cell speed in single neutrophils migrating through skin and liver capillaries. Consistent with our findings *in vitro*, we observed a positive correlation between BFAR and cell speed in both tissues. For similar speeds, the F-actin signal was higher at the cell rear in the skin capillaries (Figure 6i), which is consistent with data from vessels of varying sizes produced *in vitro* (Supplementary figure 4g).

Overall, these data demonstrate that neutrophils possess an intrinsic capacity to sense the confinement strength imposed by capillaries *in vivo* in a tissue-independent manner. We conclude that F-actin localization at the cell rear positively correlates with neutrophil migration speed in capillaries and increases in narrower channels *in vivo*.

### ROCK is essential for efficient trafficking of large number of neutrophils in a complex capillary network

Our results show that neutrophils possess the ability to adapt their cytoskeleton to varying levels of confinement in order to maintain high migration speed in straight capillaries (Figure 2). However, during their recruitment to inflamed tissues, neutrophils might encounter capillary networks of increased complexity, such as interconnected capillaries with varying diameters in the liver (supplementary figure 6c). Therefore, we hypothesized that mNeu adaptation to confinement might allow neutrophils to avoid slowing down and accumulating when facing confinement changes in such a complex microenvironment.

To evaluate whether speed maintenance is required for efficient neutrophil migration in a complex capillary network, we designed a microfabricated device of interconnected capillaries of varying diameters. This device contains four regions with an identical pattern of capillaries that differ only in their width, ranging from 3 to 6 µm (Figure 7a, supplementary Figure 7a). To mimic the massive recruitment of neutrophils during inflammation, we injected a large number of mNeu though a large peripheral channel (Figure 7a, b, supplementary Figure 7b). As a result, neutrophils could access the four regions and undergo different degrees of confinement depending on the vessel section (Figure 7c-e). We focused our analysis of neutrophil motility between 5 and 10 hours after initial entry into the device, a time window during which the average cell speed was stable in the four device areas (Supplementary Figure 7c). Notably, neutrophils exhibited similar average speed across all regions of the device, despite the increased complexity of the capillary network (Figure 7f, Supplementary Figures 7a–e, supplementary movie S21). Likewise, their capacity to explore the capillary network was independent of capillary width, as indicated by a similar mean square displacement (MSD) (Figure 7g). This data indicates that neutrophils can explore complex networks of capillaries composed of vessels of different diameters with a similar efficacy *in vitro*. To block neutrophil adaptation to confinement, we treated mNeu with the ROCK inhibitor Y-27632. Similarly to results in straight vessels, this treatment progressively reduced neutrophil speed and MSD as the confinement strength increased (Figure 7h, i; Supplementary Figures 7f–h). When comparing the MSD relative to the 6 µm-wide capillaries, Y-27632–treated cells showed a strong reduction in the 3 µm-wide capillary network (Figure 7j, supplementary movie S21). This result indicated that ROCK inhibition hampers the capacity of neutrophils to explore this network of capillaries, with a preferential effect in the smaller vessels, in agreement with the results obtained in straight tubes.

**Figure 7:**
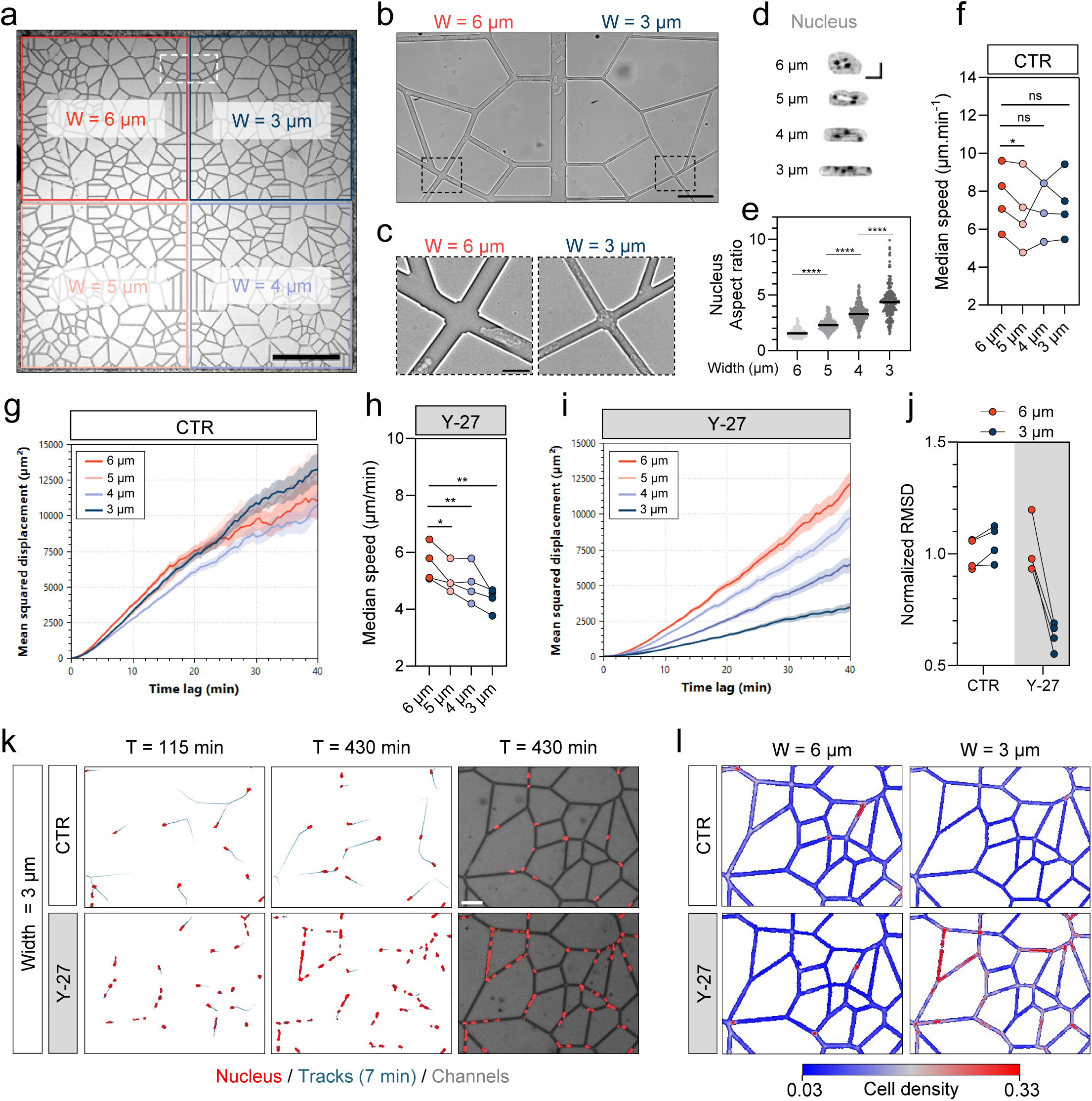
ROCK is essential for efficient trafficking of large number of neutrophils in a complex capillary network. **a**. Network of interconnected channels of different widths (6 / 5 / 4 / 3 µm) with a central channel of 20 µm with, with a height of 4 µm. Channel width regions are indicated with colored squares. mNeu are plated in a outside channel of 50 µm height and entered spontaneously in the network. Scale bar 500 µm. **b**. Magnification of the indicated region in Fig 7a. Scale bar 50 µm. **c**. Magnification regions indicated in Fig 7b. Scale bar 10 µm. **d**. Examples of mNeu nucleus (stained with Hoechst) in the indicated channel size. **e**. Nuclear AR of mNeu. From one experiment, 6 µm, 361 cells; 5 µm, 364 cells; 4 µm, 288 cells; 3 µm, 275 cells. Mann Whitney test, ****, P < 0.0001. **f**. Median speed of mNeu at late timepoints (5-10 h). From 4 different replicates of 2 independent experiments, with a minimum of 1300 cell trajectories per replicate (per channel size). One-tailed paired t test, ns, not significant. * P < 0.05. **g**. Mean squared displacement of control mNeu in 6 / 5 / 4 / 3 µm channel width. h. Median speed of Y-27632 (25 µM) treated mNeu at late timepoints (5-10 h). From 4 different replicates of 2 independent experiments, with a minimum of 890 cell trajectories per replicate (per channel size). One-tailed paired t test, ns, not significant, * P < 0.05; ** P < 0.01. **i**. Mean squared displacement of Y-27632 (25 µM) treated mNeu in 6 / 5 / 4 / 3 µm channel width. **j**. Root mean square displacement (RMSD, time lag: 30 min) of control and Y-27632 treated mNeu normalized to control in 6 µm width. **k**. Displacement of control (top) and Y-27632 treated (bottom) mNeu in the 3 µm width network, at 115- and 430-min. Nucleus are indicated in red, and tracks of 7 min are indicated in blue. Scale bar 50 µm. **l**. Cell density mapping in the 6 µm (left) and 3 µm (right) width network of control (top) and Y-27632 treated (bottom) mNeu. Color mapping represents the percentage of time that a position is covered by a cell (stained with Hoechst) for a total of 10 hours.

At the macroscopic level, ROCK inhibition in 3 µm-wide capillaries resulted in “trains” of neutrophils that blocked the vessels (Figure 7k, supplementary movie S21). Notably, this phenotype was not observed in control cells, nor in wider capillary networks, even for Y-27632-treated cells (Figure 7l). Taken together, these results show that efficient trafficking of neutrophils in a complex network of confining capillaries relies on ROCK activity. This data unveils a cellular mechanism controlling neutrophils navigation specifically into small capillaries.

## DISCUSSION

The function of immune cells is dependent on their migration capacity, and neutrophils are particularly adept at racing through tissues to reach inflamed sites. Numerous studies have demonstrated their ability to infiltrate even the narrowest spaces. However, the prevailing paradigm is that excessive confinement should eventually impair their migration. Here, we report a counterintuitive finding: neutrophils maintain the same speed regardless of whether they are migrating through large or small capillaries, both *in vivo* and *in vitro*. We further discovered that this is not because they are insensitive to physical constraints. On the contrary, neutrophils adapt their actomyosin cytoskeleton to confinement strength in just seconds. This adaptation mechanism relies on confinement-induced MyoII activation, which enables neutrophils to enhance contractility and maintain fast migration in small capillaries. Our findings reveal that MyoII activation is controlled by the ROCK pathway, independently of calcium signaling. Additionally, we show that neutrophils’ ability to maintain speed in narrow spaces operates independently of the confining material, whether endothelial cells *in vivo* or PDMS channels with various coatings *in vitro*. This suggests an autonomous mechanism for sensing confinement strength and adjusting migration speed. We propose that this adaptation helps neutrophils avoid jamming when they arrive in large numbers in tissues with small capillaries of varying diameters, a critical process for proper blood drainage and tissue function.

### Neutrophil migration in blood capillaries of various sizes

Our results show that the diameter of blood capillaries ranged from 2 µm to 10 µm, based on data from mouse skin and liver, which is consistent with previous reports (*6,7,14,40*) . In larger vessels, the average neutrophil size was 7.3 µm in diameter. Notably, neutrophil shape varied from fully round to highly elongated in smaller capillaries. Among the neutrophils, 40% exhibited active migration behavior within the capillaries, sometimes moving against the bloodstream. This observation aligns with reports indicating increased neutrophil adhesion and active migration in vessels across various tissues during inflammation (*7–9,41,42*). We also found that neutrophil migration was not impaired by the confinement of small capillaries in both the skin and liver, despite differences in basal migration speed. These findings suggest that neutrophils have an intrinsic ability to sense confinement strength in capillaries, independent of the local biochemical microenvironment. Such a mechanism could help prevent capillary obstruction due to neutrophil accumulation, regardless of the adhesion molecules expressed by endothelial cells in each tissue (*41*). The speed of neutrophil migration *in vivo* may also be influenced by the presence of inflammatory molecules. Several chemokines have been shown to facilitate neutrophil movement through restricted microenvironments (*22,43*). The chemotactic effect on neutrophils might be temporary due to desensitization of chemokine receptors (*44,45*). However, cell-intrinsic adaptation to constrained spaces could ensure neutrophil migration in narrow vessels, regardless of the inflammatory state. Notably, vascular dysfunction resulting from neutrophil stalling in capillaries has been documented in pathological conditions such as Alzheimer’s disease, ischemic stroke, or epilepsy (*14,16,17*). Further investigation is needed to determine whether this phenomenon is related to altered migratory behavior of neutrophils.

### Neutrophil migration into artificial capillaries of various sizes

The *in vitro* data presented in this study provide detailed insights into neutrophil migratory behavior in capillaries of varying sizes, independent of blood flow or chemokine signaling. In this system, the tubular structure of the vessels dictates cell shape while imposing fluid resistance and friction, which increases as confinement strength rises (*10,13*). Our research highlights that neutrophils can migrate efficiently even under extreme confinement, adopting an elongated shape with a length ten times their width. In leukocytes, the forces driving cell migration are primarily regulated by the organization of the actin cytoskeleton at the protrusive front and the contractile rear (*33*). Actomyosin plasticity has been shown to regulate cell migration in microchannels, whether through cell activation or genetic or pharmacological perturbations (*46–48*). Here, we demonstrate that in neutrophils, actomyosin organization adapts to the microenvironmental confinement strength in under a minute. This results in neutrophils reaching the same speed after a brief slowdown when facing abrupt narrowing. Furthermore, we show that neutrophil instantaneous speed positively correlates with higher F-actin enrichment at the cell rear for a given cross-sectional area. This observation aligns with previous studies linking leukocyte speed to actomyosin localization at the cell rear (*33,46,49–52*). In confined microenvironments, cells adopt a migration mode where front actin is depleted to allow pressure-driven protrusion (*27,53,54*). In neutrophils migrating through narrow capillaries, F-actin resembles the actomyosin network self-organization observed in blebs under confinement (*27,53*). Blebs are stabilized by increased contractility at their neck through MyoII activation, leading to F-actin depletion at the front and sustained migration (*27,53*). Consistent with these findings, we show that MyoII inhibition predominantly affects neutrophil migration in narrow channels, corroborating studies in other cell types (*25–27,55,56*).

### Matching propulsive and resisting forces to maintain migration speed

The adaptation of actomyosin to produce more force under confining conditions has been documented across various cell types and organisms, even those distantly related to animal cells (e.g., choanoflagellates) (*34–36,57*). However, whether this adaptation allows cells to maintain a constant speed was previously unexplored. In this study, we found that neutrophils exhibited a unique ability to maintain their speed, a feature not observed in other leukocytes. To better understand this, we examined single cells, plotting the degree of actin enrichment at their rear (BFAR) versus their speed for each channel size (Supplementary figure 4j). As expected, for any channel size, more actin at the rear corresponds to faster cells, suggesting that BFAR can be used as a proxy for the propelling force exerted by the cell. When comparing various channel sizes, we observed that cells with a given BFAR move more slowly in narrower channels, suggesting that more propulsion force is needed to reach a given speed in these channels. Since the relationship between BFAR and speed is typically linear, it is tempting to interpret this with the simplest physical principle: Newton’s law at low Reynolds number (without inertia). It states that propulsion forces and forces resisting migration should balance each other. Resisting forces can be described as effective friction, which increases linearly with speed (ρV), and BFAR, as a proxy for the propulsion force (Fp), which leads to the conclusion that BFAR∝Fp=ρV. This explains the increasing slope for smaller channels, corresponding to a larger friction ρ. This suggests that neutrophils are able to match the increase in friction in smaller channels with a proportional increase in propulsion force. To the best of our knowledge, no simple physical explanation exists for how propulsion force could perfectly adapt to match increased friction. We speculate that this is due to a sensing mechanism fine-tuning myosin activity based on resisting migration forces. Such specific fine-tuning, not seen in other leukocytes, is likely essential for proper neutrophil function.

### Mechanism of modulation of MyoII activation to match resisting forces

Our study reveals that MyoII activation occurs almost instantly (relative to the imaging timescale, meaning within less than a minute) when a cell enters a confined region of a capillary, and this activation is maintained during migration under confinement. This aligns with observations of cortical redistribution of MyoII during neutrophil extravasation *in vivo* (*58*). Similarly, during NK cell migration through collagen matrices, ROCK-dependent MyoII contractile bursts have been observed before full cell deformation when migrating through narrow pores (*59*). This suggests an autonomous MyoII response to deformation that may rapidly occur in leukocytes facing confinement during migration. In line with this, MyoII activation upon uniaxial confinement has been documented in various cell types (*34–36,57*). In these contexts, MyoII activity increases seconds to minutes after cell confinement, in either a calcium-dependent or calcium-independent manner, depending on the cell type (*34–36,57*).

Several mechanisms have been proposed to explain shape-dependent activation of MyoII. The most direct effect could be purely geometrical, as proposed for the modulation of small GTPases activated by GEFs localized at the plasma membrane. Making the cell flatter or more elongated increases the surface-to-volume ratio, bringing cytoplasmic GTPases closer to their activators at the plasma membrane (*60*). Such a mechanism could be used by migrating leukocytes under strong confinement. Narrower spaces might also favor increased concentrations of autocrine factors, such as leukotriene B4 (LTB4), which is required for neutrophil extravasation induced by self-production and sensing of this molecule (*58*). However, such a mechanism is unlikely in this case, given the second-time-scale activation of contractility upon entering a narrow channel and the strict correlation with changes in channel width, not consistent with changes in extracellular concentration of secreted factors.

In addition to these geometric effects, shape changes can also induce mechanical stresses due to large strains on various cellular structures. Actomyosin networks have been proposed to display strain-induced activation. However, this is unlikely in our case because cells move and deform spontaneously, as opposed to fast deformation from externally applied forces (as in the case of forced flow, compression or stretching) and because we found that upstream ROCK regulation is required for activation in narrow channels. We also provided strong evidence that calcium, the classic deformation-sensing molecule, although it increases inside cells in narrower channels, is not required for speed maintenance. This rules out all calcium-dependent pathways, including the cPLA2-dependent nuclear ruler pathway (*34,35*). The increase in intracellular calcium is likely due to membrane tension, which is associated with the opening of channels on plasma or intracellular membranes (*38*). Although calcium itself does not seem involved in the fast adaptation of actomyosin in neutrophils, the rise of intracellular calcium suggests that membrane tension also increases in cells entering narrow channels. Recent studies have shown that membrane tension can serve as an upstream signal for the integration of mechanical stress across the whole cell (*61*). Indeed, membrane tension can propagate across the cell surface, coordinating protrusions and contractions in neutrophils (*62–64*). De Belly and colleagues suggested that local increases in membrane tension at one part of the cell could result in RhoA activation at the opposite side. This signaling can occur within seconds via the mechanosensitive mTORC2 pathway (*64*). Similarly, neutrophil entry into narrow capillaries could increase membrane tension at the leading edge, causing RhoA activation at the rear. In agreement with this, we observed rapid cortical MyoII enrichment at the cell rear in neutrophils entering small capillaries, detected even before full cell deformation (Figure 5e). Sustained high membrane tension in small vessels would then contribute to sustained RhoA and MyoII activation, maintaining propulsion force. The maintenance of the speed of cells migrating in continuously narrowing channels (Figure 3j) suggests that the adaptation mechanism does not rely on a threshold but rather on a continuous increase of the propulsive force and Myosin II activity in response to increased resisting forces.

A plausible working model based on these observations might involve a feedback mechanism, where increased resistance in narrower capillaries raises membrane tension, possibly due to an increased friction on the walls as the cell elongates. This, in turn, would enhance RhoA and myosin activity, effectively adjusting the propulsive force. But what ensures the proportionality of these processes, required to maintain a constant speed, remains an open question, and might be the result of fine tuning by selection pressure. This would suggest an important function of speed maintenance in neutrophils.

### Functional relevance of speed maintenance in leukocytes

We observed that tissues contain capillaries small enough to be fully obstructed by a single neutrophil (Figure 1a). When a local infection occurs, there is rapid recruitment of neutrophils to these capillaries (Figure 1a). In such a context, any systematic slowdown caused by capillary geometry would likely lead to jamming. To test this hypothesis, we conducted preliminary experiments in networks of interconnected capillaries with varying diameters, filled with large numbers of neutrophils. We considered both control and ROCK inhibition, where neutrophils could not adapt to the channel diameter and systematically slowed down. While control cells never jammed, ROCK-inhibited cells rapidly accumulated and formed immobile “trains” in channels of small diameter (Figure 7k, l). We conclude that neutrophils’ unique ability to fine-tune their propulsion force to match the resisting forces imposed by their environment is crucial to avoiding jamming in small capillaries with varying dimensions. Further studies are required to support this conclusion in a more physiological context.

Interestingly, other leukocytes tested did not display this capacity to maintain their average speed when facing narrowing capillaries. Previous studies have shown that other leukocytes, such as dendritic cells, adapt their myosin activity to confinement, which is necessary for efficient migration in narrow spaces. However, most leukocytes do not maintain speed in smaller capillaries. In fact, their speed can be reduced by half under such conditions (Figure 3k, l). This cannot be explained by larger cell size, as T cells and NK cells—cells with similar sizes to neutrophils—do not maintain their speed when confined in capillaries. It has been reported that, unlike most cells, neutrophils are not limited in their migration by their nucleus, while other cells are (*12,65,66*). This is attributed to the highly deformable morphology and mechanical properties of neutrophil nuclei. While this factor may contribute to our findings, it does not fully explain our results. Our data show that neutrophils need to adapt their actomyosin to maintain their speed and that narrower channels produce more resisting forces, even for these highly compliant cells. We also previously showed that larger cells with stiffer nuclei, such as activated dendritic cells, can maintain their speed in capillaries of varying confinement levels *in vitro* (*56*), suggesting that neutrophils share this capacity with these cells. Together, these observations indicate that the maintenance of speed in leukocytes migrating in capillaries may be linked to their specific function and activation state (*55,67*).

In summary, our study reveals that neutrophils can continuously adapt their migration machinery to changes in confinement strength imposed by capillaries of varying diameters. This ability likely contributes to the efficiency of neutrophil trafficking in vessels during inflammation, minimizing the risk of vascular dysfunction.

## MATERIALS AND METHODS

### Mice

Male and female mice on a C57BL/6J (B6) genetic background were used at age ranging from 8 to 16 weeks. These mice were either wild-type (C53BBL/6J inbred strain, obtained from Charles Rivers) or genetically altered to express the Lifeact-EGP peptide (kind gift from M. Sixt) (*68*) or the Myosin IIA-GFP fusion protein (kind gift from R. S. Adelstein) (*32*). These mouse strains did not exhibit a harmful phenotype thus their breeding and maintenance are not subjected to a project evaluation. Mice were housed under specific pathogen–free conditions within the animal research facility of the Institut Curie, in compliance with the European and French National Regulation for the Protection of Vertebrate Animals used for Experimental and other Scientific Purposes (Directive 2010/63; French Decree 2013-118). Mice were euthanized by cervical dislocation by appropriately qualified and trained individuals.

### Intravital experiments

For intravital imaging, experiments were performed at Institut Pasteur in agreement with guidelines established by the French and European regulations for the care and use of laboratory animals and approved by the Institut Pasteur Committee on Animal Welfare (CETEA) under the protocol code CETEA 230080. C57BL/6 male and female mice were used at age ranging from 6 to 14 weeks and were housed under standard conditions (light 07.00–19.00 h; temperature 22 ± 1 °C; humidity 50 ± 10%) within the animal facility of the Institut Pasteur.

30-min prior to intravital imaging, mice were injected subcutaneously with buprenorphine (0.05 mg kg-1) and anesthetized by spontaneous inhalation of isoflurane in 100% oxygen (induction: 4%; maintenance: 1.5% at 0.3 L min−1). Mouse hydration and body temperature were respectively maintained by intraperitoneal injection of 200 μl 0.9% saline solution every hour and with 37 °C-heating pad. Labelling antibodies were injected intravenously 15 min prior to imaging, 5µg of anti-Ly6G (clone 1A8) conjugated with Alexa-Fluor 488, 561 or 647 to label circulating neutrophils and 10 µg of CD31 (clone 390) conjugated with Alexa-Fluor 488, 561 or 647 to label the vasculature. Mice were imaged for up to 4 hours.

For infected dorsal back skin observations, mouse preparation was conducted as previously described(69). Using Nanofill Syringe (World Precision Instruments), 3 intradermal injections of 1 µl of 10^8^ CFU ml−1 bacterial suspension of the wild-type *N. meningitidis* strain were made(7). For liver observations, organ preparation was conducted as previously described (*70*). Mice were treated with an intraperitoneal injection of 1 mg/kg purified LPS from E. coli 2-hours prior to intravital microscopy. For LifeAct-GFP imaging in dorsal back skin and liver, purified bone-marrow LifeAct-GFP expressing neutrophils (100 µl, 7.5×10^6^ cells) were injected intravenously 3-hour prior imaging.

Intravital imaging was performed using a Leica DM6 FS upright microscope equipped with a motorized stage. The microscope is fitted with HCX PL Fluotar 5x/0.15, HC Fluotar ×25/0.95 and HC PL APO ×40/ 1.10 objectives lens (Leica), mounted on an optical table to minimize vibration and totally enclosed in a biosafety cabinet (Noroit). The microscope is coupled to a Yokogawa CSU-W1 confocal head modified with Borealis technology (Andor). Four laser excitation wavelengths (488, 561, 642, and 730 nm) were used and visualized with the appropriate band-pass and long-pass filters. Fluorescence signals were detected using 2 sCMOS 2048 × 2048 pixels cameras (Orca Flash v2+, Hamamatsu) allowing simultaneous dual colors imaging. Metamorph acquisition software (Molecular devices) was used to drive the confocal microscope.

### Mouse neutrophil isolation and culture

A single-cell suspension was prepared from mouse femur and tibia’s bone marrow. Mouse neutrophils were isolated by immunomagnetic negative selection using the Mojosort neutrophil isolation kit (BioLegend, #480058) and according to the manufacturer’s instructions. Expression of surface markers of neutrophils were analyzed by flow cytometry to assess the sample purity (supplementary figure 8). Neutrophils enrichment was assessed by flow cytometry analysis of Ly6G and CD11b expression.

After isolation, neutrophils were cultured overnight at a concentration of 1×10^6^ cells/mL in RPMI 1640 medium supplemented with fetal bovine serum (10 %), penicillin-streptomycin (100 U/mL) and recombinant granulocyte-macrophage colony-stimulating factor (GM-CSF, 50 ng/mL, Peprotech, #315-03-20UG).

### Bone-marrow derived dendritic cells *in vitro* differentiation

The BMDCs were obtained by differentiation of bone marrow precursors for 10 days in IMDM-Glutamax medium supplemented with fetal bovine serum (10%), penicillin-streptomycin solution (100 U/ml), and β-mercaptoethanol (50 μM) and granulocyte-macrophage colony-stimulating factor (GM-CSF)-containing supernatant (50 ng/mL) obtained from transfected J558 cells, as previously described (Vargas, NCB). At day 10, semi-adherent cells were collected by carefully flushing with media after discarding the floating cells. Full differentiation was assessed by flow cytometry analysis of CD11c and CD80 expression.

### Mouse T cell isolation, activation and expansion

Whole spleens were harvested from mice then dissociated in PBS into single-cell suspensions by passing the splenocytes through a sterile 70 μm mesh nylon strainers (Falcon, 352350). Naïve CD4+ T cells from mouse splenocytes were isolated by immunomagnetic negative selection, using the EasySep Mouse naïve CD4+ T Cell Isolation Kit (StemCell, # 19765). Isolated T cells were activated using monoclonal antibodies against the ε chain of mouse CD3 (subunit of the TCR complex) and the mouse CD28 co-stimulatory molecule for 24 hours and expanded in DMEM medium containing fetal bovine serum (10%), non-essential amino acids (1%), penicillin-streptomycin (100 U/mL), sodium pyruvate (1 mM), Beta-mercapto-ethanol (50 µM) and recombinant interleukine-2 (100U/mL, Preprotech, #212-12). T cells were used after 6 days of culture.

### Mouse NK cell isolation and expansion

Fresh whole spleens were harvested from mice then dissociated in PBS into single-cell suspensions by passing the splenocytes through a sterile 70 μm mesh nylon strainers (Falcon, 352350). Subsequently, NK cells were enriched by negative selection with magnetic nanobeads using the MojoSort Mouse NK Cell Isolation Kit (Biolegend, cat. # 480050) according to the manufacturer’s instructions. NK enrichment was assessed by flow cytometry analysis of NK1.1+/CD3-expressing cells. NK cells were cultured in 12-well flat-bottomed plates (Nest, cat. # 712001) at a concentration of 1×10^6^ cells/mL in culture media containing RPMI 1640 medium supplemented with FBS (10 %), non-essential amino acids (1 %), HEPES buffer, β-mercaptoethanol (55 µM), and recombinant interleukine-15 (7 nM, Biolegend, cat. # 566302). NK cells were used after 4-5 days of culture.

### Photolithography

All microdevices used for this work were designed and prepared as follows. First, chrome photomasks were fabricated by JD photo Data (UK) from our designs with specific dimensions (negative pattern). Next, using photolithography technique, SI wafers were photopatterned with an SU-8 negative photoresist. The obtained Si wafers were subsequently silanized and a thick layer of PDMS (RTV615, Neyco) was poured over the wafer and cured at 65°C for at least 2h. PDMS was peeled off the Si wafer and used to prepare a replica mold using epoxy.

### Microchannel preparation

Microchannels experiments were performed as previously described (*46*). Briefly, A 10-to-1 mixture of PDMS (RTV615, Neyco) was poured into the custom-made epoxy molds. PDMS crosslinking was obtained after 2 hours at 65°C. PDMS chips were peeled off from the epoxy mold using a blade, cut at the edges, then 2- or 3-mm holes were drilled in the PDMS chamber in a region near the entry of microchannels, as a reservoir for cell loading. PDMS chamber and 35-mm glass-bottom dish (FD35-100, WPI) were cleaned then plasma activated before being bound to each other. The binding was left to strengthen in a 65 °C oven for 15 min.

For surface functionalization, the PDMS surfaces were plasma activated during 1 min and incubated with the coating solution at RT during 30 min. For most experiments, microchannels were coated with fibronectin (10 μg/ml, Sigma-Aldrich, #F1141). Alternatively, when specified, microchannels were coated with collagen (10µg/mL, Advanced BioMatrix, #5005), ICAM-1 (10µg/ml), pLL-PEG (1mg/ml) or Pluronic-F127 (50mg/ml). After the incubation step, PDMS chambers were washed three times with PBS then incubated with complete medium (containing drugs when specified) for 2 hours at 37 °C and 5% CO2 before cell loading.

### Widefield time-lapse microscopy

For velocity measurements and actin imaging, migrating cells were recorded for 6 hours with a DMi8 inverted microscope (Leica) at 37 ◦C with 5% CO2 atmosphere and a ×20 dry objective (NA 0.40 phase). One image every 30s during 6 h was recorded. The widefield microscope was controlled by MetaMorph software.

### Spinning-disk time-lapse microscopy

For MyosinII imaging, migrating cells were recorded using a spinning-disk confocal microscope with a Yokogawa CSU-X1 spinning-disc head on a DMI-8 Leica inverted microscope equipped with a Hamamatsu OrcaFlash 4.0 Camera, a NanoScanZ piezo focusing stage (Prior Scientific) and a motorized scanning stage (Marzhauser). One slice with focus on the middle cell cross section was recorded every 5 or 10 seconds for 1 hour. The spinning-disk microscope was controlled by MetaMorph software (Molecular Devices) and equipped with an incubation chamber that maintained the temperature at 37°C and CO2 concentration at 5%.

### Image processing

All images were exported from the Metamorph acquisition software (Molecular Devices) as .tiff files and edited using Fiji. Stacks were registered using Stackreg plugin or using SIFT multichannel plugin.

### Image analysis

Three single-cell parameters (cell AR, cell speed, and F-actin intracellular distribution) were analyzed using in-house ImageJ macros. Briefly, kymographs for each microchannel (in each field of view) were generated using a semi-automated ImageJ macro. Individual cells migrating within the microchannels were segmented on each kymograph using a pixel classification algorithm (ilastik and Labkit, applied to the LifeAct-EGFP or MyoII-EGFP signal) to obtain a mask of the entire cell. The cell AR, used as a measure of cell deformation/elongation (Viridis LUT), was computed with ImageJ as the ratio of the major axis to the minor axis of the Feret ellipse approximating the cell. Instantaneous cell speed was determined by measuring the displacement of the cell centroid along the microchannel axis between two frames. The back-to-front actin ratio (BFAR) was calculated to evaluate intracellular F-actin distribution. The BFAR corresponds to the ratio of the mean F-actin fluorescence intensity measured in the rear half of the cell mask to that measured in the front half. A BFAR greater than 1 indicates an enrichment of F-actin at the rear of the cell, while a BFAR less than 1 indicates an enrichment of F-actin at the front of the cell.

### Spatial analysis of single-cell parameters

The relative speed of cells migrating in microchannels with sharp transitions was calculated for each cell as the ratio of its instantaneous velocity to its average velocity in the first section of the microchannel. In microchannels with successive transitions, the average velocity of each cell was calculated in the first 6 µm section. In microchannels with a gradual narrowing, the relative speed of each cell was computed as the ratio of its instantaneous velocity to its average velocity.

To assess the evolution of BFAR, AR and speed along the x axis of the microchannels, we partitioned the x axis into fixed-size bins. Then, the average across cells in each bin was calculated. To avoid sampling bias and underrepresentation of the fastest cells, we partitioned the x axis into bins with a length equal to the largest cell displacement measured in the experiment. By doing so, each bin contains at least one measure for each cell. When more than one measure for a cell was recorded in a specific bin, these measures were averaged before the averaging across the cell population. Relative speed data and BFAR data were log2 transformed prior to the averaging across the cell population.

The crossing time between two microchannels sections was calculated as the time between the first cell contact with the narrowing section of the microchannel, and the time at which the whole cell was in the confined section.

### Lifeact-GFP density maps

Lifeact-GFP density maps were generated as described(46). Briefly, individual neutrophils migrating in microchannels were cropped using the ImageJ software. To map the Lifeact-GFP signal, images of all timepoints of each individual cell were aligned in a single column. Cell size normalization was applied to each time point according to the mean cell size. To obtain the mean Lifeact-GFP distribution of each cell, an average projection was performed on the t-stack. All cell average projections were then size-normalized and averaged to obtain the mean Lifeact-GFP distribution of each condition.

### Immunofluorescence

PDMS device (coated with fibronectin) with a transition from 6 x 4 µm (large) to 3 x 4 µm (narrow) channels were stick on 18 mm diameter glass coverslips. Neutrophils were prepared as mentioned above, plated in microchannels, and incubated for 2-3 hours at 37°C to let them migrate in large and narrow channels. Cells were fixed with 4% paraformaldehyde for 20 min at 37°C and washed with PBS. PDMS device was gently removed by first unsticking PDMS from coverslips with razor blades and then pulling vertically with tweezers. Cells were permeabilized with 0.1 % Triton X-100 for 10 min at room temperature, washed and blocked with PBS-BSA 3% overnight at 4°C. Cells were with stained with rabbit anti-pMLC (Cell Signaling Technology, 3674) for 24h at 4°C, washed, stained with a goat anti-rabbit AF647 (Invitrogen, A32733) for 1 hour at RT, and washed again. Cells were stained for 5 min with DAPI (Invitrogen, D1306) and the coverslip was mounted on a glass slide with Fluoromount G (Molecular probes). Images were taken with the spinning-disc confocal microscope, with a 40X objective. Z stacks of 20 slices every 0.5 µm were taken. Image analysis was performed using ImageJ/Fiji software (NIH, https://rsb.info.nih.gov/ij/index.html). ROIs of single cells in 6 x 4 µm or 3 x 4 µm channels were cropped. Z max projection of Phalloidin signal was used for cell area segmentation, and P-MLC mean intensity was measured within cell area for every slices. For images presented in (Figure 5a), z max projection of P-MLC signal was represented, with cell contours (from cell segmentation).

## Acknowledgments

we thank Ana-Maria Lennon Duménil (Institut Curie, Paris, France) and Fernando Sepulveda (Institut Imagine, Paris, France) for sharing mice stocks. We thank Matthieu Bernard for helping with raw data processing from Metamorph. We thank the animal facility at the Institut Curie, and the microscopy facility of IPGG for equipment and technical assistance.

## Author contributions

M.D. performed and analyzed most experiments; M.B. analyzed data of *in vitro* experiments; P.N. performed the *in vivo* experiments; M.M. wrote ImageJ macros for image processing and analysis; L.B. performed experiments with immature dendritic cells; L.W. performed photolithography; T.J. helped with image analysis; E.T. performed FACS analysis; O.B. performed TL experiments; A.D. helped with image processing and analysis; S.G. helped with analysis; M.G.G. performed NK experiments; E.T. and R.A. did initial experiments with dendritic cells; G.D. supervised *in vivo* experiments; M.D., M.P., and P.V., designed the study. M.D., P.V. and M.P. wrote the manuscript. All authors contributed to discussions and the final version of the manuscript.

---

**Supplementary figure 1:**
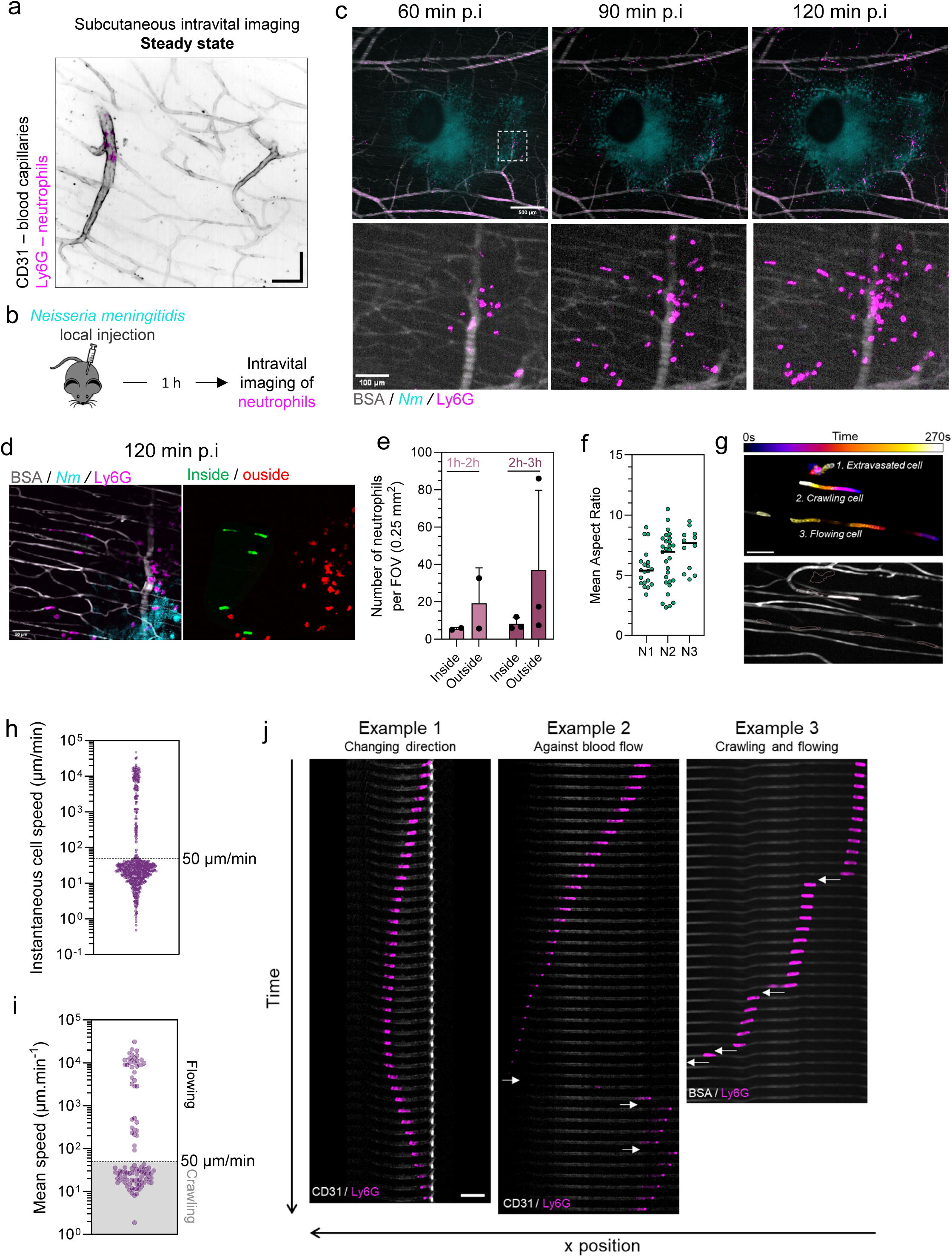
**a.** Spinning disk-confocal images of subcutaneous blood capillaries (inverted gray, fluorescent BSA) and endogenous neutrophils (magenta, Ly6G) at steady state. Scale bar: 20 µm. **b**. Experimental strategy to observe endogenous neutrophil trafficking in subcutaneous blood capillaries by spinning-disk-confocal intravital imaging. **c**. Top, 5X objective spinning disc-confocal images of subcutaneous blood capillaries (gray, fluorescent BSA), endogenous neutrophils (magenta, Ly6G), and *Neisseria meningitidis* (cyan) at 60 min, 90 min and 120 min post-infection. Scale bar: 500 µm. Bottom, magnification of white square indicated in top image. Scale bar: 100 µm. **d**. Left, 25X objective spinning disk-confocal images of subcutaneous blood capillaries (gray, fluorescent BSA), endogenous neutrophils (magenta, Ly6G), and *Neisseria meningitidis* (cyan) at 120 min post infection. Scale bar: 50 µm. Right, neutrophils located inside (green) or outside (red) capillaries. **e**. Number of neutrophils inside and outside capillaries per field of view (0.25 mm^2^) between 1h and 2h and between 2h-3h post-infection. Bars represent mean +/- SD from two (1h-2h) or three (2-3h) independent experiment. **f**. Mean cell aspect ratio of neutrophils crawling inside subcutaneous capillaries of three independent experiments. **g**. Time projection of one field of view, showing a neutrophil migrating outside capillaries (1), a neutrophil crawling inside capillary (2), and a neutrophil intermittently crawling and flowing inside capillary (3). **h**. Instantaneous trafficking speed of individual cells in subcutaneous capillaries, n = 878 timepoints of 99 cells from 3 independent experiments. **i**. Mean trafficking speed of individual cells in subcutaneous capillaries. Data are pooled from 3 independent experiments, n = 99 cells. **h**. Examples of neutrophil migratory behavior inside skin capillaries. Example 1 shows a neutrophil doing a U-turn. Example 2 shows a neutrophil migrating towards the left and being flowed toward the right three times (white arrows). Example 3 illustrates a cell crawling toward the left and being flowed in the same direction (white arrows).

**Supplementary figure 2:**
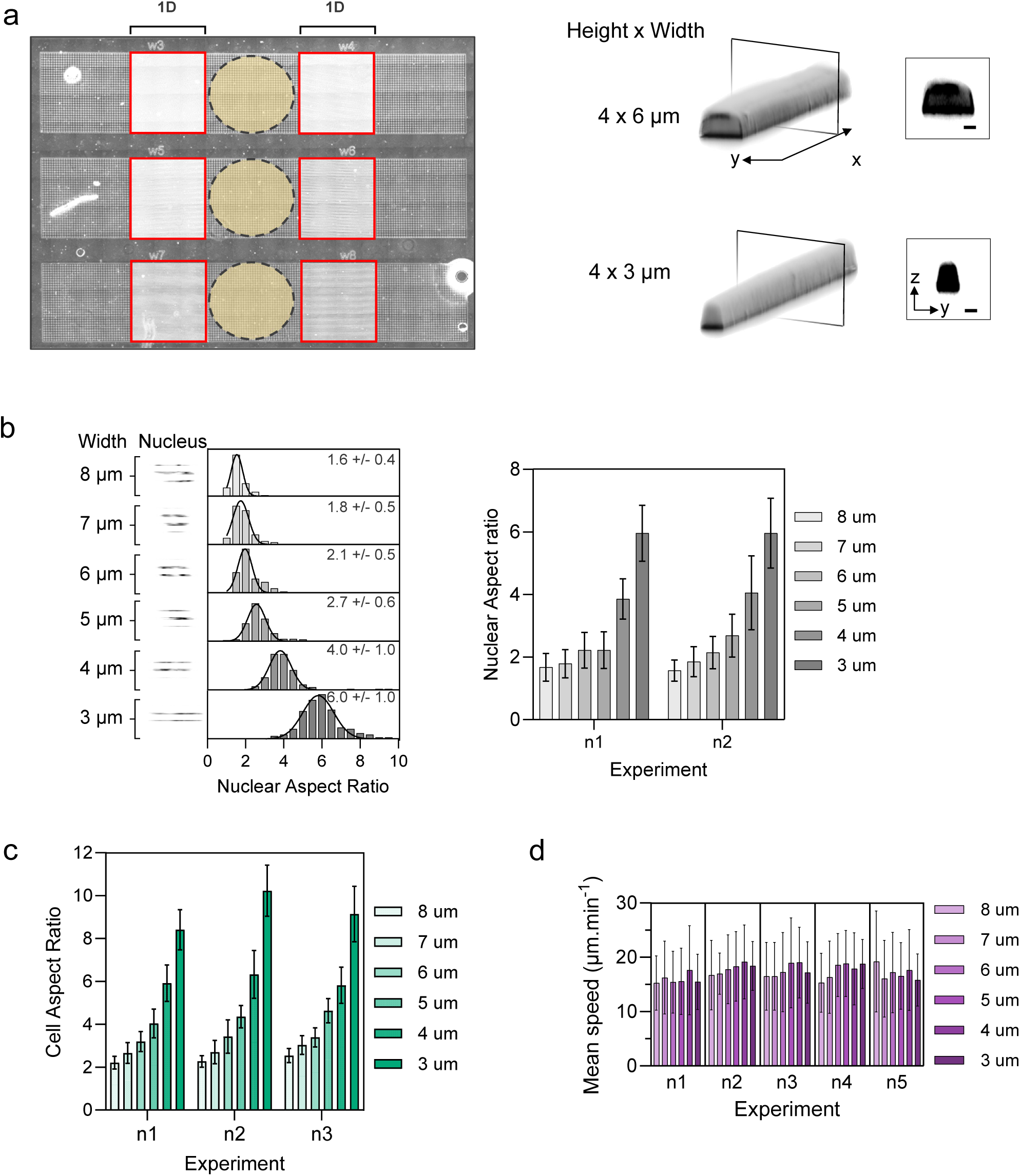
**a**. Left, image of PDMS-microfabricated device of straight microchannels. Three rows of chambers are aligned. Each row contains two microchannels regions of a specific width (indicated with red rectangles). A hole is made in the yellow region, where 100k cells were plated. Cells spontaneously migrated to the left and right and entered in microchannels. Right, 3D reconstruction of a 4 x 6 µm channel section (top) and a 4 x 3µm channel section using fluorescent dextran. **b**. Left, Distribution of mean nuclear aspect ratio, from 2 independent experiments. Median +/- SD is indicated for each channel width. 8 µm, n = 80; 7 µm, n = 86; 6 µm, n = 88; 5 µm, n = 118; 4 µm, n = 132; 3 µm, n = 137. Right, mean +/s SD of mean nuclear aspect ratio for each independent experiments. **c**. Mean cell aspect ratio +/s SD for each independent experiments (pooled in Fig 2.c). **d**. Mean cell speed +/s SD for each independent experiments (pooled in Fig 2.d).

**Supplementary figure 3:**
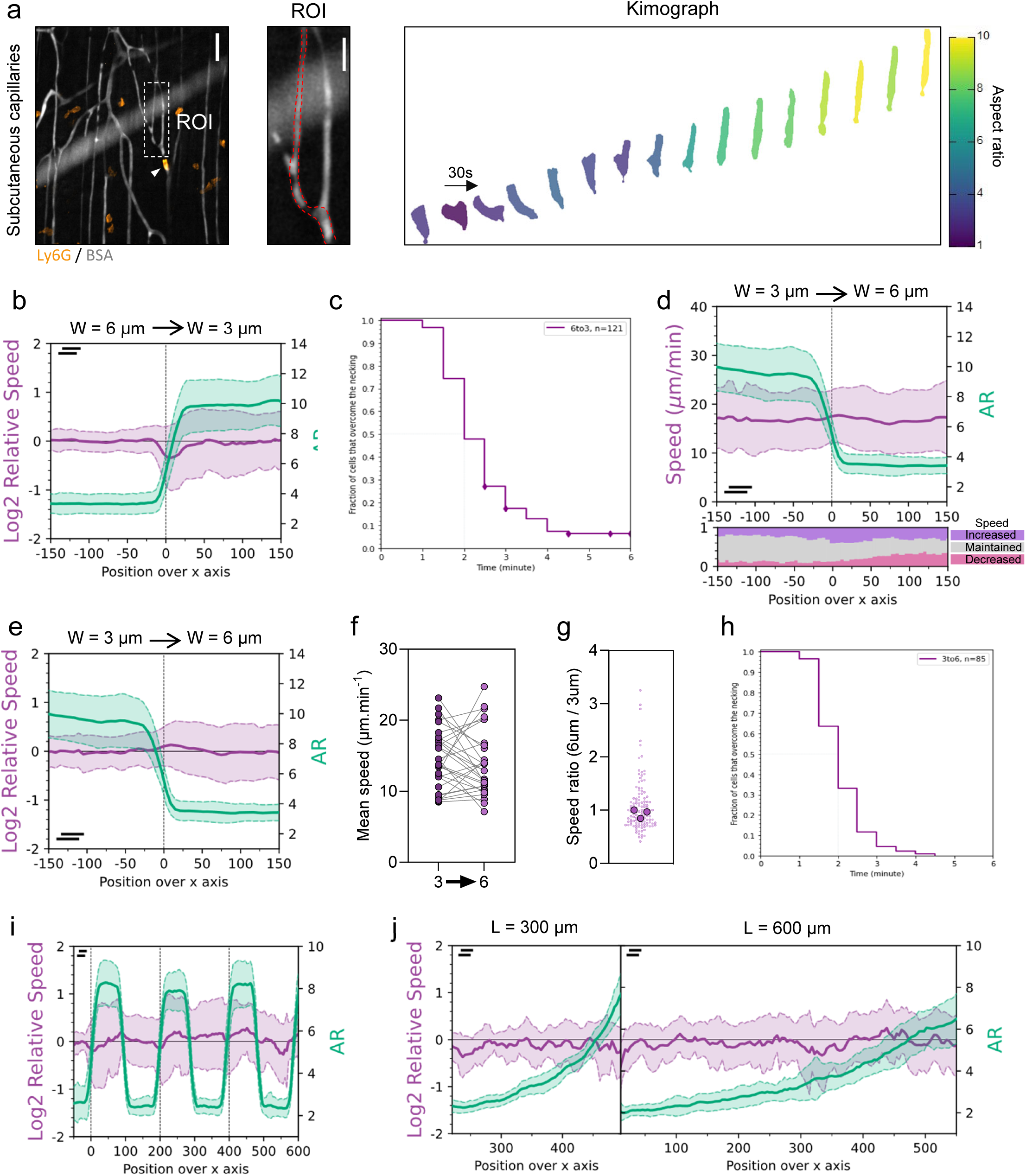
**a**: Example of a mNeu migrating in a subcutaneous capillary exhibiting gradual narrowing. Right, kymograph showing mNeu displacement with color-coded cell aspect ratio. **b**: Relative speed (Log2) and cell aspect ratio over the x position in a size-transition microchannel (corresponding to figure 3b). Position 0 corresponds to the necking. Data are one representative of 5 experiments, N = 122 cells. **c**: Fraction of mNeu crossing the constriction (necking) from 6 to 3 µm-wide channel according to the time. At 2 min, 50 % of mNeu have entirely entered in the 3 µm-wide section. Data are one representative of 5 experiments, N = 121 cells. **d**: Top, evolution of cell speed and cell aspect ratio over the x position in a size-transition microchannel. Position 0 corresponds to the enlargement from 3 to 6 µm width. Bottom, evolution of the fraction of cells displaying a speed increase, maintenance or decrease over the x axis, compared with the respective mean speed in the 3 µm-wide section. Data are one representative of 3 experiments, N = 86 cells. **e**: Relative speed (Log2) and cell aspect ratio over the x position in a size-transition microchannel (corresponding to supplementary figure 3d). **f**: Average speeds of 30 cells migrating from 3 µm to 6 µm-wide section. **g**: Speed ratio: average speed in the 6 µm-wide section normalized by the average speed in the 3 µm-wide section. Each small dot represents a cell, and bigger dots represent a median of one experiment. From 3 independent experiments, N = 119 cells. **h**: Fraction of mNeu crossing the channel enlargement from 3 to 6 µm width according to the time. At 2 min, 50 % of mNeu have entirely left the 3 µm-wide section. Data are one representative of 3 experiments, N = 85 cells. **i**: Relative speed (Log2) and cell aspect ratio over the x position in a successive size-transition microchannel (corresponding to figure 3g). **j**: Relative speed (Log2) and cell aspect ratio over the x axis of the funnel-shaped channel of 300 µm length (left) and 600 µm length (right). Corresponding to figure 3j.

**Supplementary figure 4:**
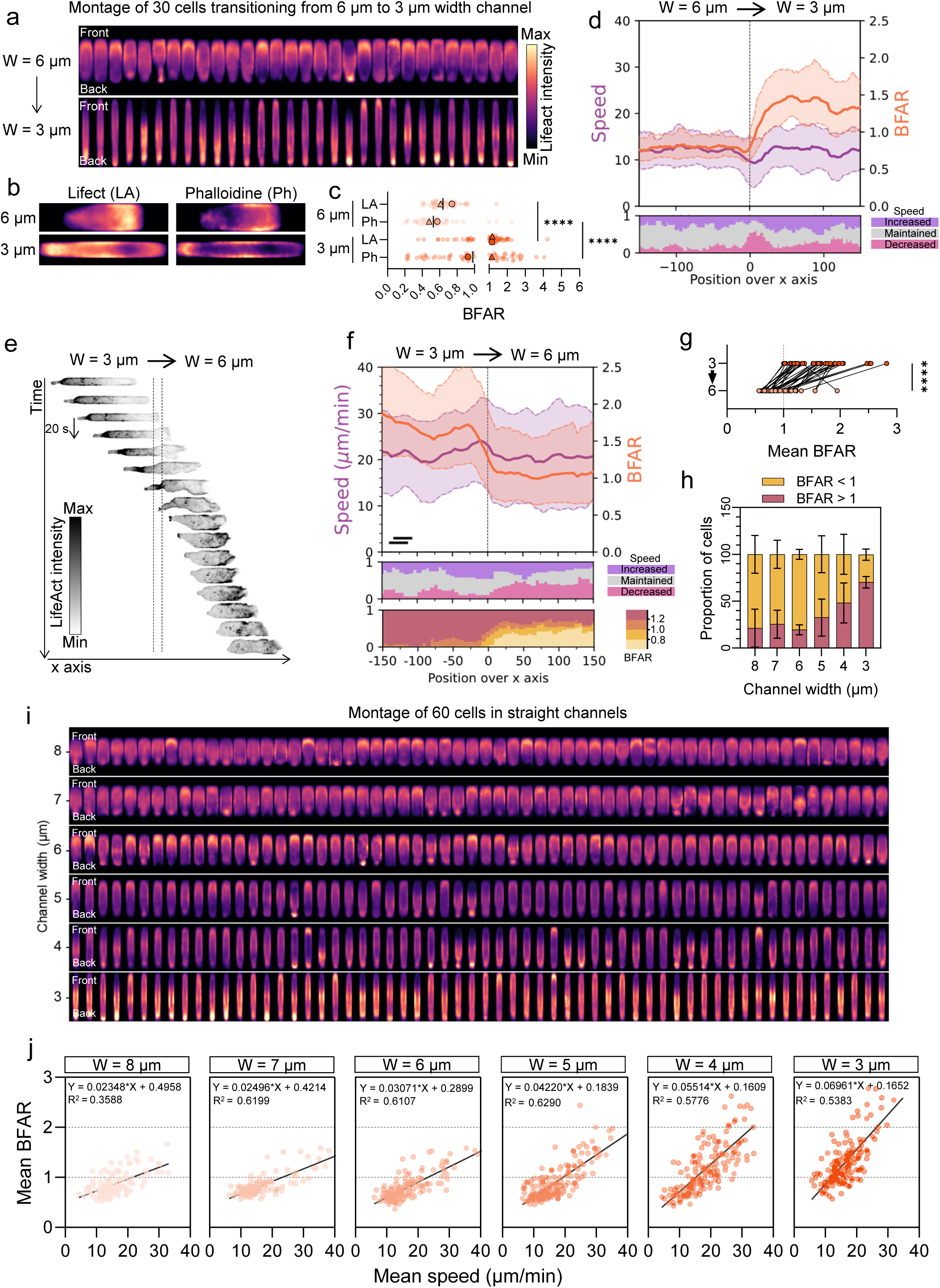
**a**: Average mapping of lifeact-EGFP of 30 mNeu migrating from 6 µm to 3 µm width channels. Each row represents a cell, in the 6 µm (top) and the 3 µm (bottom) channel. **b**: Average mapping of lifeact-EGFP (left) and phalloidine staining (right) of fixed lifeact-EGFP expressing mNeu in 6 µm (top) and 3 µm (bottom) wide channels. Data are one representative of two independent experiments. 6 µm, n = 35 cells; 3 µm, n = 49 cells. **c**: Mean BFAR measured with lifeact-EGFP and Phalloidine signal of fixed lifeact-EGFP expressing mNeu in 6 µm and 3 µm wide channels. Data are pooled from two independent experiments. 6 µm, n = 44 cells; 3 µm, n = 61 cells. Two-tailed Mann Whitney test, **** P < 0.0001. **d**: Top, BFAR and cell speed over the x position in a size-transition microchannel. Position 0 corresponds to the necking. Bottom, evolution of the fraction of cells displaying a speed increase, maintenance or decrease over the x axis, compared with the respective mean speed in the 6 µm wide section. Data are one representative of 4 experiments, n = 50 cells. **e**: Kimograph of a Lifeact-EGFP expressing neutrophil migrating from 3 µm to 6 µm wide channel. Each row represents a timepoint (20 seconds). Lifeact-EGFP intensity is represented in inverted gray color. **f**: Top, BFAR and cell speed over the x position in a size-transition microchannel. Position 0 corresponds to the transition from 3 µm to 6 µm wide section. Middle, evolution of the fraction of cells displaying a speed increase, maintenance or decrease over the x axis, compared with the respective mean speed in the 6 µm wide section. Bottom, fraction of cells over the x axis, with BFAR values between indicated BFAR ranges. Data are one representative of 3 experiments, n = 25 cells. **g**: Mean BFAR of individual neutrophils migrating from the 3 µm to the 6 µm wide section. Data are pooled from three independent experiments, n = 40 cells. Two-tailed Wilcoxon test (paired), **** P < 0.0001. **h**: Proportion of mNeu with a mean BFAR value below or above one, in straight microchannels of different widths. Data are the mean +/- SD of 4 independent experiments. 8 µm, n = 216; 7 µm, n = 210; 6 µm, n = 257; 5 µm, n = 289; 4 µm, n = 271; 3 µm, n = 253 cells. **i**: Average mapping of lifeact-EGFP of 60 cells (20 cells of 3 independent experiments) migrating in straight channels of different widths. **j**: Relation between BFAR and cell speed of neutrophils migrating in straight channels of different widths. Mean values for individual cells are represented. Simple linear regression is applied with indicated slope and R squared. Data are pooled from 3 independent experiments. 8 µm, n = 132; 7 µm, n = 120; 6 µm, n = 170; 5 µm, n = 187; 4 µm, n = 169; 3 µm, n = 163 cells.

**Supplementary figure 5:**
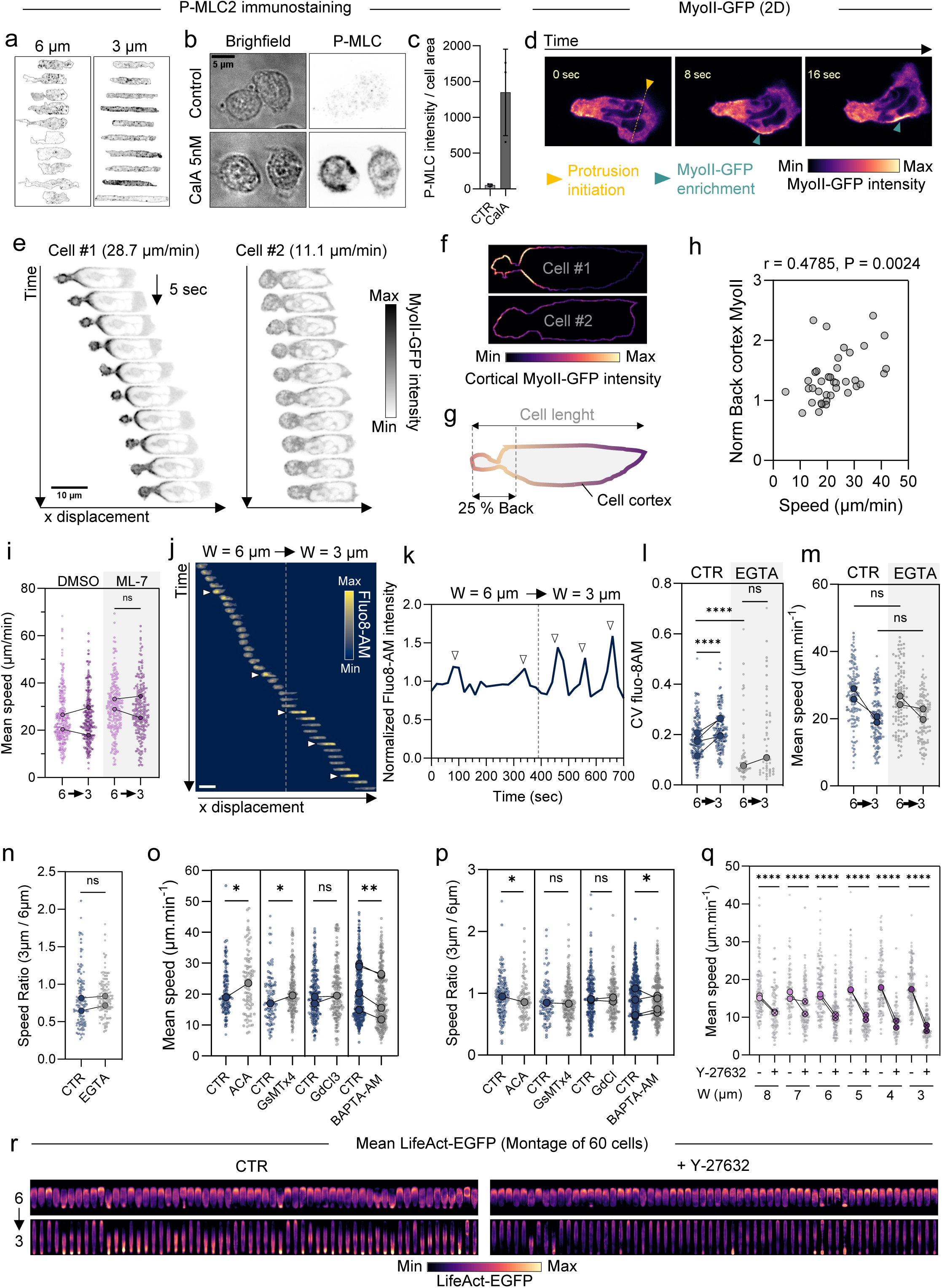
**a**. Immunostaining of Phospho-Myosin Light Chain 2 (P-MLC2) of fixed neutrophils in 6 µm and 3 µm-wide microchannels. Ten cells are represented. P-MLC2 signal is in inverted gray. **b**. Immunostaining of Phospho-Myosin Light Chain 2 (P-MLC2) of two control and Calyculin A treated neutrophils. **c**. P-PMLC2 levels of control and Calyculin A treated neutrophils. **d**. Example of a MyoII-GFP neutrophil changing direction during migration in 2D confined. **e**. Kimographs of a fast (Cell #1, left) and slow (Cell #2, right) MyoII-GFP expressing mNeu. MyoII-GFP signal is represented in inverted gray. **f**. Cortical MyoII-GFP signal of cell #1 and cell #2 (one timepoint represented). **g**. Illustration of back cortical MyoII-GFP measurement. **h**. Correlation between mean cell speed and back cortical MyoII signal (normalized to total MyoII signal). Spearman correlation coefficient = 0.48, P = 0.002 (two tailed). **i**. Average speeds of cells migrating from 6 µm to 3 µm-wide section, treated or not with ML-7. Data are pooled from 2 independent experiments. DMSO, n = 257; ML-7, n = 217. Mann Whitney test, ns, not significant. **j**. kymograph showing the evolution of cytosolic calcium signaling (Fluo8-AM) of a mNeu migrating in a size-transition channel. White arrows indicate calcium pulses. Vertical dotted line indicates the narrowing. **k**. Evolution of Fluo8-AM levels the migrating mNeu represented in j. Vertical line represents the time of cell passage in the 3 µm-wide section. **l**. Coefficient variation (Standard deviation / Mean) of Fluo8-AM intensity of control and EGTA (1 mM) treated cells. Data are pooled from 4 (CTR) and 1 (EGTA) independent experiments. CTR, n = 165 cells; EGTA, n = 48 cells. Mann Whitney test; ns, not significant; **** P < 0.0001. **m**. Average speeds of cells migrating from 6 µm to 3 µm-wide section, treated or not with EGTA (1 mM). Data are pooled from 2 independent experiments. Big connected dots represent the median of each experiment. Mann Whitney test; ns, not significant. **n**. Speed ratio of mNeu treated or not with EGTA (1 mM). Each light dot represents a cell, and connected dark dot represents medians of each experiment. Data are pooled from 2 independent experiments. CTR, n = 129; EGTA, n = 113. Mann Whitney test; ns, not significant. **o**. Average speeds of cells migrating in 6 µm-wide section, treated or not with ACA, GsMTx4, GdCl, or BAPTA-AM. Data are pooled from 1 (ACA), 1 (GsMTx4), 2 (GdCL3) and 4 (BAPTA-AM) independent experiments. DMSO (ACA), n = 147 cells; ACA, n = 95; CTR (GsMTx4), n = 100, GsMTx4, n = 193; CTR (GdCl3), n = 232, GdCl3, n = 182; CTR (BAPTA-AM), n = 472, BAPTA-AM, n = 299 cells. Mann Whitney test; ns, not significant; * P < 0.05; ** P < 0.01. **p**. Speed ratio of mNeu treated or not with ACA, GsMTx4, GdCl, or BAPTA-AM. Data are pooled from 1 (ACA), 1 (GsMTx4), 2 (GdCL3) and 4 (BAPTA-AM) independent experiments. Connected big dots represent the median of each experiment. DMSO (ACA), n = 147 cells; ACA, n = 95; CTR (GsMTx4), n = 100, GsMTx4, n = 193; CTR (GdCl3), n = 232, GdCl3, n = 182; CTR (BAPTA-AM), n = 472, BAPTA-AM, n = 299 cells. Mann Whitney test; ns, not significant; * P < 0.05. **q**. Average speeds of mNeu in straight channels of different widths (W), treated (+Y) or not (-Y) with Y-27632. Data are pooled from 2 independent experiments. 8 µm: - Y = 122, +Y = 97; 7 µm: -Y = 80, +Y = 136; 6 µm: -Y = 121, +Y = 134; 5 µm: -Y = 118, +Y = 119; 4 µm: -Y = 131, +Y = 94; 3 µm: -Y = 163, +Y = 107. Mann Whitney test; **** P < 0.0001. **r**. Average Lifeact-EGFP intensity mapping of control and Y-27632 treated cells in size-transition channels. 60 cells are represented from 3 independent experiments (20 cells per experiment).

**Supplementary figure 6:**
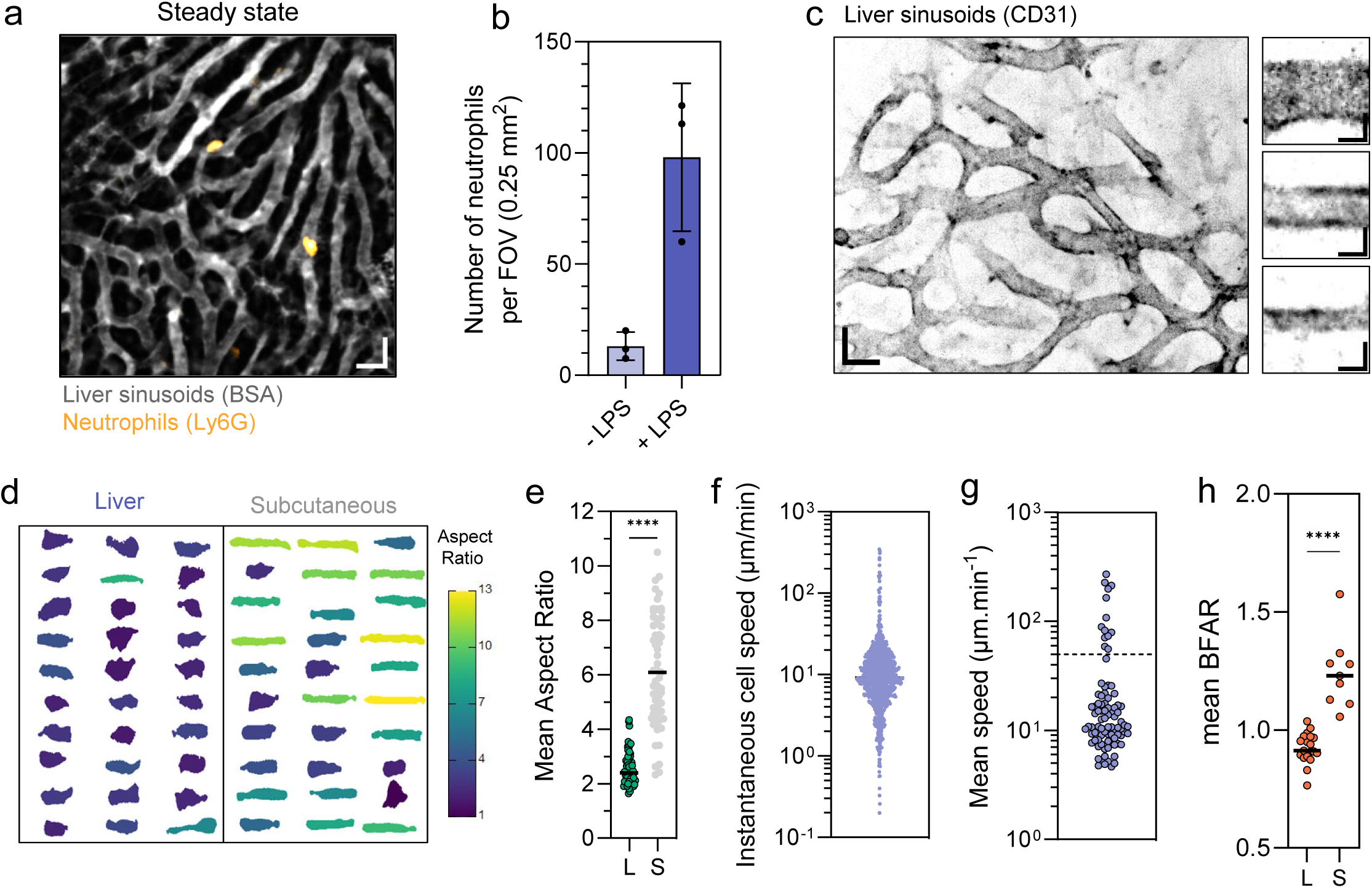
**a**: Spinning disk-confocal images of liver sinusoids (gray, fluorescent BSA) and endogenous neutrophils (orange, Ly6G) at steady state. Scale bar: 20 µm. **b**: Quantification of neutrophils number at steady state and 2 hours after LPS endotoxemia. Each dot represents the mean of 3 independent experiments. **c**. Left, spinning disk-confocal images of liver sinusoids (inverted gray, CD31). Scale bar 20 µm. Right, magnifications of 3 sinusoid segments. Scale bar 5 µm. **d**. Montage of 30 cells trafficking in liver sinusoids (left) and subcutaneous capillaries (right), pseudo-colored by the value of their AR. Montage of subcutaneous condition is the same than figure 1e. **e**. Mean cell AR of neutrophils in liver sinudoids (L) and subcutaneous capillaries (S). Data of subcutaneous condition are the same than figure 1g. Data are pooled from 3 independent experiments. L, n = 69 cells; S, n = 61 cells. Mann Whitney test, **** P < 0.0001. **f**. Instantaneous trafficking speed of individual cells in liver sinusoids, n = 1098 timepoints of 87 cells from 3 independent experiments. **g**. Mean trafficking speed of individual cells in liver sinusoids. Data are pooled from 3 independent experiments, n = 87 cells. **h**. Mean BFAR of mNeu expressing lifeact-EGFP mNeu migrating in liver sinusoids (L) or subcutaneous capillaries (S). Data are pooled from 2 (L) and 1 experiment (S). L, n = 19 cells; S, n = 9 cells. Mann Whitney test, **** P < 0.0001.

**Supplementary figure 7:**
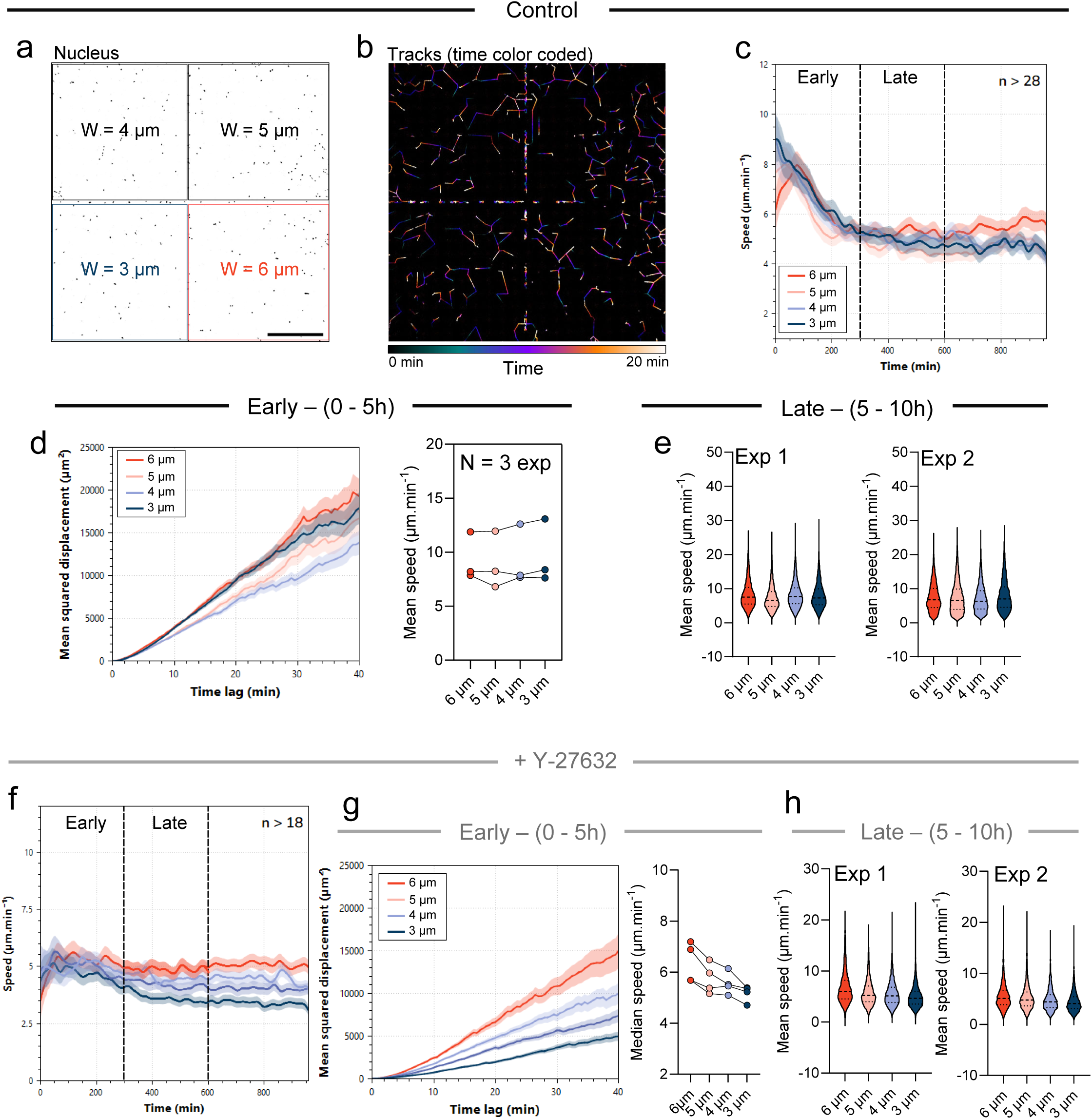
**a**. Image showing nucleus (inverted gray) of mNeu in the device of channels network. Widths of channels are indicated on the image. Scale bar 500 µm. **b**. Image showing 20 min tracks of mNeu nucleus in the device. **c**. Evolution of mean speed of non-treated mNeu over time. Standard error is represented in light color. Early (0-5 h) and late (5 – 10 h) time windows correspond to the time after mNeu entry in networks. Data are representative of 4 replicates from 2 independent experiments, with a total of more than 2200 cell trajectories per channel width (minimum of 28 trajectories at any timepoint). **d**. Migratory behaviors of mNeu in the networks of channels of varying widths, at early timepoints (0 – 5h). Left, mean squared displacement of mNeu. Data are one representative of 6 replicates from 3 independent experiments. Right, Mean speed of mNeu population. Data represent the mean of 3 independent experiments, with a minimum of 1300 trajectories per channel width per experiment. **e**. Mean speed of mNeu trajectories in the networks of channels of varying widths, at late timepoints (5 – 10 h). From two independent experiments (left and right), with a minimum of 4000 trajectories per channel width per experiment. **f**. Evolution of mean speed over time of Y-27632 treated mNeu. Standard error is represented in light color. Early (0-5 h) and late (5 – 10 h) time windows correspond to the time after mNeu entry in networks. Data are representative of 4 replicates from 2 independent experiments, with a total of more than 2200 cell trajectories per channel width (minimum of 18 trajectories at any timepoint). **g**. Migratory behaviors of Y-27632 treated mNeu in the networks of channels of varying widths, at early timepoints (0 – 5h). Left, mean squared displacement of mNeu. Data are one representative of 4 replicates from 2 independent experiments. Right, Mean speed of mNeu population. Data represent the mean of 4 replicates from 2 independent experiments, with a minimum of 500 trajectories per channel width per replicate. **h**. Mean speed of Y-27632 treated mNeu trajectories in the networks of channels of varying widths, at late timepoints (5 – 10 h). From two independent experiments (left and right), with a minimum of 1600 trajectories per channel width per experiment.

**Supplementary figure 8:**
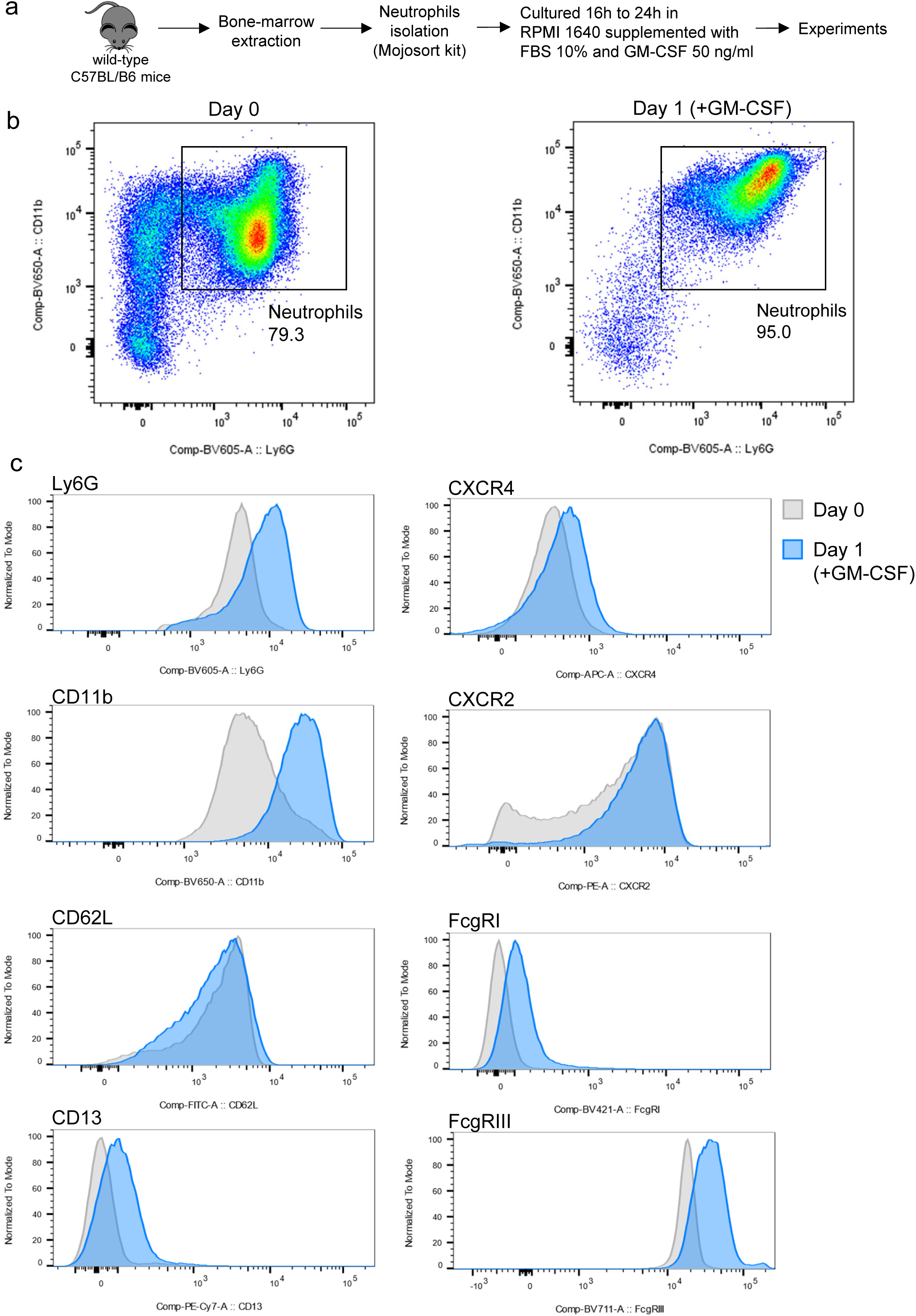
**a**. Experimental protocol for mouse bone marrow neutrophil preparation. **b**. FACS quantification of bone marrow neutrophils. Ly6G fluorescence is analyzed in CD11b+ cells at day 0 after isolation (left) and at day 1 (in the presence of GM-CSF, right). **c**. Flow cytometry analysis of indicated protein surface expression at day 0 and day 1 (GM-CSF).

